# Transcriptomic profiling of epigenetic regulators and metabolic reprogramming in human cholangiocarcinoma

**DOI:** 10.64898/2025.12.10.693445

**Authors:** Amaya Lopez-Pascual, Jasmin Elurbide, Emiliana Valbuena-Goiricelaya, M Ujue Latasa, Elena Anaya, Elena Adan-Villaescusa, Borja Castelló-Uribe, Luz A. Martínez-Pérez, Iker Uriarte, Maria Arechederra, Sergio Ciordia, Fernando J Corrales, Sona Frankova, Eva Sticova, Ondrej Fabian, Leticia Colyn, Patricia Inacio, Robert Arnes-Benito, Juan Bayo, Meritxell Huch, Carmen Berasain, Maite G Fernández-Barrena, Matías A Avila

**Author notes:** **Correspondence:** Corresponding authors: Amaya Lopez-Pascual, PhD and Matias A Avila, PhD. Hepatology Laboratory, Solid Tumors Program, CIMA, CCUN, University of Navarra. Avda. Pio XII, n55. 31008 Pamplona, Spain. MGF-B and MAA share senior authorship.

## Abstract

**Background:** Epigenetic alterations play an increasingly recognized role in carcinogenesis and in the development of resistance to anticancer therapies. Epigenetic enzymes (writers and erasers) and effectors (readers) are largely influenced by the availability of metabolites grenerated through one-carbon metabolism (OCM), the tricarboxylic acid (TCA) cycle, and acetyl-CoA synthesis (ACS). In this study we examined the expression of epigenetic and metabolic genes to investigate their interplay in cholangiocarcinoma (CCA).

**Method:** We examined 257 epigenetic genes (EpiGs), 96 metabolic genes (MGs), and 189 rate-limiting enzymes (RLEs) in transcriptomic data from iCCA, eCCA, CCA organoids, and normal bile ducts, alongside prognostic signatures. CRISPR-Cas9 DepMap data evaluated the impact of EpiGs and MGs on cell viability. HuCCT-1 iCCA cells were exposed to hypoxia (1% O₂, 24 h) to assess EpiG responses. Transcriptomic deconvolution characterized EpiGs, MGs, and RLEs expression across four tumor microenvironment (TME) subtypes. Two mouse CCA models (Akt/TAZ, Akt/NICD) underwent RNA-seq, complemented by multi-omic profiling (transcriptomic, proteomic, metabolomic) in Akt/TAZ livers.

**Results:** Several EpiGs were upregulated in iCCA and eCCA, including writers (*DNMT1*, *EZH2*, *SUZ12*), readers (*CBX3*, *PHF20L1*, *SMARCA4*), and erasers (*HDAC1*, *HDAC3*, *KDM5C*). MGs in OCM, TCA, and ACS pathways were dysregulated (up: *GART*, *IDH2*, *TYMS*; down: *ALDH1L1*, *MAT1A*, *SHMT1*). Integrated analyses identified 27 EpiGs and 8 MGs whose overexpression predicted poor survival. Subsets of EpiGs, MGs, and RLEs were linked to proliferative, high-recurrence iCCA subclasses. CRISPR screens highlighted 50 EpiGs and 23 MGs essential for CCA viability. Tumor microenvironment (TME) analyses revealed distinct immune-stromal subclasses with coherent epigenetic-metabolic signatures. In CCA cells, hypoxia induced epigenetic programs that mirrored those in CCA patients. Transcriptomic analyses in human and multi-omic analyses (transcriptomic, metabolomic, and proteomic studies) in mouse CCA livers highlighted rewiring of nucleotide, one-carbon, lipid, and mitochondrial pathways, with evidence of metabolic-epigenetic crosstalk.

**Conclusion:** EpiGs and MGs are markedly altered in both human and experimental CCA, with several changes particularly enriched in aggressive molecular subclasses associated with poor prognosis. We observed substantial rewiring of epigenetic cofactor-related MG expression in CCAs. Functional assays validated new targets among EpiGs (e.g. *CBX3*, *CHD4*, *DEK*, *SMARCA4*, and *TRIM28*) and MGs (*TYMS* and *IDH2*) in CCA.

## 1 Introduction

Cholangiocarcinoma (CCA) is the second most frequent primary liver cancer and a very aggressive epithelial cell malignancy (Valle et al., 2021). While there are geographical differences regarding the underlying etiological and risk factors, the incidence and mortality rates associated with CCA are globally rising (Brindley et al., 2021; Pascale et al., 2023). According to its anatomical origin CCA can be classified as intrahepatic CCA (iCCA) when it occurs in the biliary tree within the liver parenchyma, and perihilar CCA (pCCA) or distal CCA (dCCA), when the lesions emerge outside the liver, and in some studies these tumors are collectively referred to as extrahepatic CCA (eCCA) (Brindley et al., 2021; Valle et al., 2021; Pascale et al., 2023). CCA is a lethal disease with a 5-year overall survival ranging from 7-20% (Banales et al., 2020; Ilyas et al., 2023). The reasons for this very poor prognosis are multifarious and may include a late diagnosis (Macias et al., 2022), when curative resection or liver transplantation are no longer possible (Brindley et al., 2021; Valle et al., 2021), and also the intrinsic resistance of CCA cells to cytotoxic therapy (Brindley et al., 2021; Valle et al., 2021). CCAs are molecularly heterogeneous tumors. Indeed, a variety of genetic alterations like mutations in genes such as *TP53*, *ARID1A*, *KRAS*, *SMAD4*, *BAP1*, *IDH1*, *PI3KCA*, *BRAF*, translocations in the *FGFR2* gene, and *ERBB2* amplifications, that represent potent oncogenic drivers, have been identified (Brindley et al., 2021; Ilyas et al., 2023). Noteworthy, some of these alterations show differential prevalence depending on the anatomical subsite of the tumors, like *FGFR2* fusions and *IDH1* mutations which are more frequent in iCCA, and *KRAS* mutations and *ERBB2* amplifications in pCCAs and dCCAs (Brindley et al., 2021; Ilyas et al., 2023). Interestingly, certain genetic aberrations such as *FGFR2* fusions and *IDH1* mutations are amenable to pharmacological targeting, and there are approved drugs showing promising results in recent clinical trials (Alvaro et al., 2023; Ilyas et al., 2023). However, the prevalence of actionable mutations among different CCA types is variable (Valle et al., 2021; Ilyas et al., 2023), and the emergence of acquired resistance to FGFR inhibitors in patients harboring *FGFR2* fusions is beginning to be documented (DiPeri et al., 2023). The complex and evolving molecular profile of CCA may underlie its high resistance to chemotherapy and also towards targeted agents (Marin et al., 2018; O’Rourke et al., 2023). Importantly, it is becoming evident that genetic alterations cannot account for all tumor characteristics, including the variable speed at which the disease progresses along its natural history, or the differential response to chemotherapy (Chaisaingmongkol et al., 2017; O’Rourke et al., 2023). While acquired somatic mutations certainly impact the transcriptional profile and thus the behavior of CCA cancer cells (Ahn et al., 2019; Dong et al., 2022), the role of epigenetic alterations in the remodeling of CCA transcriptome and tumor progression is increasingly recognized (O’Rourke et al., 2019; Manzano-Núñez et al., 2023; Zhong et al., 2023).

Different pathways are involved in chromatin dynamics and epigenetic gene regulation, including DNA methylation, ATP-dependent nucleosome remodeling complexes, post-translational modifications (PTMs) of histones, and a variety of non-coding RNAs, acting in an intricate crosstalk (Du et al., 2015; O’Rourke et al., 2018; Fernández-Barrena et al., 2020; Claveria-Cabello et al., 2021; Zhao et al., 2021). Histones PTMs modulate chromatin compaction and control the recruitment of remodeling complexes and transcription factors (Morgan and Shilatifard, 2020). These covalent PTMs are reversible marks introduced, removed and recognized by a broad set of proteins that according to these functions can be classified as epigenetic writers, erasers and readers (Biswas and Rao, 2018; Fernández-Barrena et al., 2020). Dysregulation of epigenetic pathways can occur through different mechanisms, including genetic mutations in epigenetic modifiers and the aberrant expression and activity of key chromatin remodelers. Some of these alterations have been reported in CCA, such as mutations in chromatin remodelers (*e.g*. *ARID1A*), histone methyltransferases (*e.g. KMT2C*, *KMT2D*), histone demethylases (*e.g. KDM6B*), histone acetyl readers (*e.g. PBRM1*); or the overexpression of histone and DNA methyltransferases (*EZH2*, *EHMT2*, *DNMT1*), histone deacetylases (*HDAC3*) and the epigenetic scaffold *UHRF1* (Jusakul et al., 2017; O’Rourke et al., 2019; Colyn et al., 2021; Bao et al., 2022; Zhong et al., 2023). The identification of DNA methylation alterations leading to the silencing of tumor suppressor genes and the enhancement of spontaneous mutagenic events, as well as the recognition of genome-wide DNA methylation profiles associated with differential CCA patients’ prognosis, further support the involvement of epigenetic dysregulation in CCA pathogenesis (Jusakul et al., 2017; O’Rourke et al., 2019; Colyn et al., 2021; Goeppert et al., 2022; Chen et al., 2023; Dragomir et al., 2023). Importantly, for epigenetic writers and erasers, as well as for the enzymes that introduce and remove methyl groups in DNA, key cellular metabolites are needed as substrates or can behave as enzymatic inhibitors. It has become evident that fluctuations in their cellular levels can have an impact on epigenetic PTMs. Therefore, the interplay between the intracellular pools of these metabolites and the epigenetic machinery adds another layer of complexity to the regulation of gene expression and its derangement in cancer cells, which undergo extensive metabolic rewiring (Li et al., 2018; Pajares and Pérez-Sala, 2018; Satriano et al., 2019; Zhang et al., 2019; Boon, 2021; Huo et al., 2021; Raggi et al., 2021, 2022). Because certain metabolic enzymes catalyze rate-determining steps that control overall metabolic flux, their deregulation provides a direct readout of the metabolic constraints shaping cofactor availability, thereby influencing epigenetic activity in CCA (Ally et al., 2017; Boon, 2021; Huo et al., 2021).

Contrary to genetic mutations, epigenetic mechanisms such as DNA and histones covalent modifications are highly flexible and dynamic, involving reversible enzymatic reactions and specific protein-protein interactions. These features make epigenetic mechanisms amenable to pharmacological intervention, and great interest has been placed in the development of the so-called epigenetic drugs, or epidrugs, mainly for the treatment of neoplastic diseases (Bates, 2020). Several epigenetic therapies have already been tested, demonstrating beneficial effects in preclinical models of cholangiocarcinoma (Xiang et al., 2014; Jung et al., 2017; Lu et al., 2019; Colyn et al., 2021; Xu et al., 2022; Zhang et al., 2022; Chen et al., 2024). However, despite promising preclinical results, the clinical translation of epidrugs has so far achieved only limited success. This has been attributed in part to greater toxicity than expected and also to low efficacy (Feehley et al., 2023). A better understanding of the critical epigenetic alterations driving cancer may help in the selection of more efficient and less toxic therapeutic approaches, as well as their potential combinations with cytotoxic agents and immune-based therapies (Sarantis et al., 2021; Huang et al., 2022; Akce et al., 2023).

Mouse models of CCA provide a controlled setting to dissect disease mechanisms and therapeutic responses, and among the various approaches available, transposon-based systems are widely used, with TAZ/Akt and NICD1/Akt combinations representing two of the most established strategies to induce cholangiocarcinogenesis in mice (Banales et al., 2020; Cigliano et al., 2022; Ma et al., 2024). In parallel to in vivo systems, patient-derived tumoroids and non-malignant biliary organoids have become powerful tools to model cholangiocarcinogenesis and therapeutic response, as demonstrated by the liver cancer-derived organoid platform established by Broutier et al. (Broutier et al., 2017), which enables faithful recapitulation of human CCA biology and supports high-throughput drug screening.

In this study, taking advantage of publicly available data, as well as CCA tumoroids and CCA mosue models, we performed a comprehensive transcriptomic analysis of epigenetic modifiers and metabolic enzymes related to epigenetic processes in CCA tissues from different cohorts of patients. In addition, we also explored the dependency of human CCA cancer cell lines on the expression of epigenetic modifiers through the analysis of genetic screening data (Meyers et al., 2017; Behan et al., 2019; Krill-Burger et al., 2023) and their sensitivity to selected small-molecule epigenetic inhibitors.

## 2 Methods

### 2.1 Selection of epigenetic and metabolic genes

Epigenetic genes (EpiGs) were selected from the literature (Biswas and Rao, 2018), EpiFactors (Marakulina et al., 2023), and ChromoHub (Liu et al., 2012) databases essentially as previously described (Herranz et al., 2023) and were manually curated, excluding those with no experimental evidence of their functional activity in repositories (GeneCards, PubMed, and Uniprot) (**Table 1 and Supplementary Table 1**). In those EpiGs with more than one biochemical activity, categories were established prioritizing writer and eraser functions over reader or cofactor activities (Herranz et al., 2023). Metabolic genes functionally linked to epigenetic regulation (MGs), including those participating in folate and methionine metabolism within the one-carbon metabolism (OCM) pathway, the tricarboxylic acid cycle (TCA), and acetyl-CoA synthesis (ACS), are shown in **Supplementary Table 2**. Additionally, an extended analysis of metabolic reprogramming was performed using 92 metabolic pathways downloaded from the KEGG resource (Kanehisa et al., 2014). Genes from each of the 57 pathways are reported in **Supplementary Table 3**. Rate-limiting enzymes (RLEs) were retrieved from RLEdb (Zhao et al., 2009), containing 189 genes encoding human RLEs. To ensure updated annotation, each enzyme entry was manually reviewed using the BRENDA (https://www.brenda-enzymes.org/) and ExPASy Enzyme (https://enzyme.expasy.org/) databases. This manual curation was used to verify the current Enzyme Commission (EC) number, associated metabolic pathway, and functional classification, ensuring accurate integration with our transcriptomic datasets (**Supplementary Table 4**).

**Table 1.**
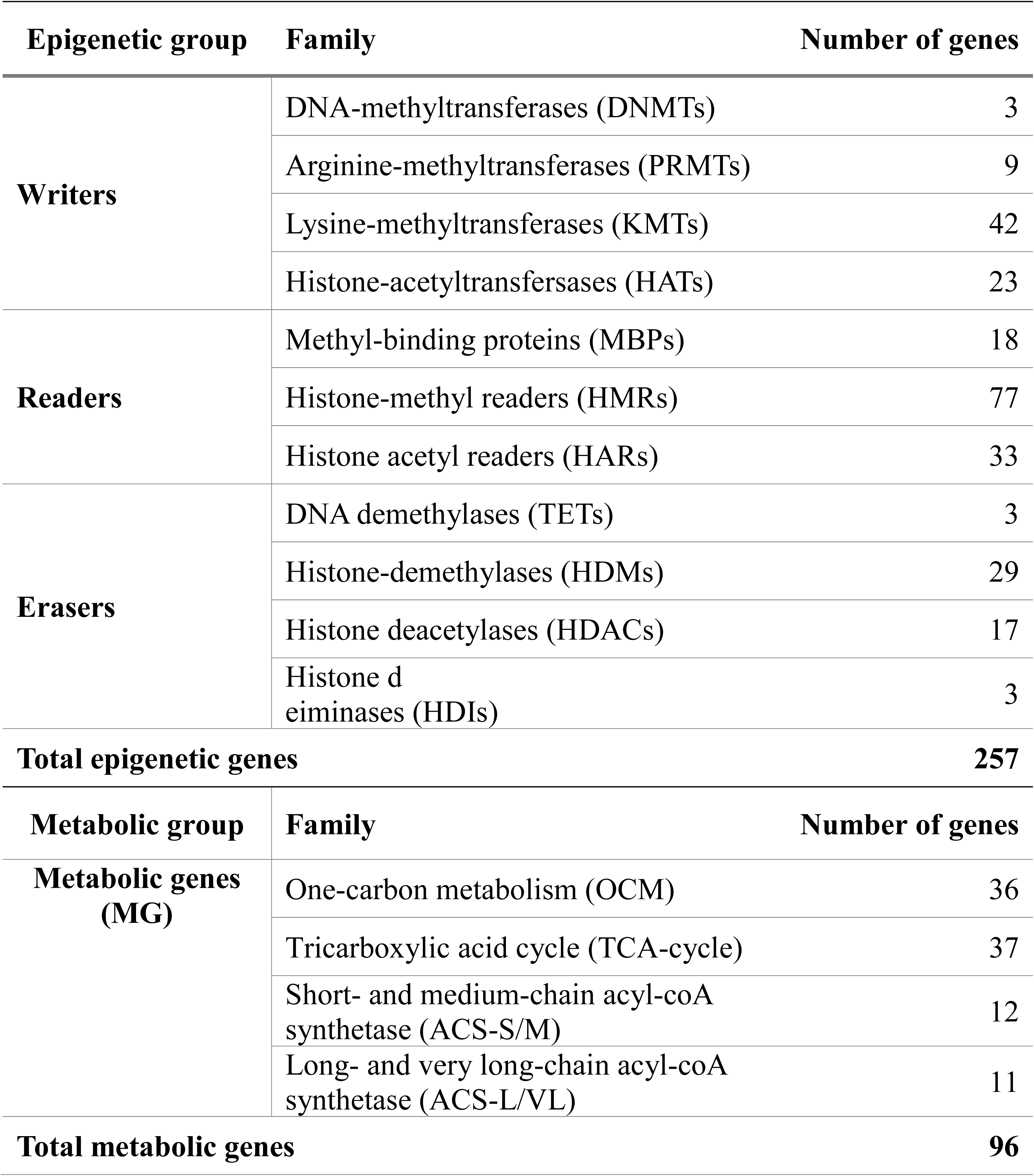
Epigenetic and metabolic genes classified by families.

### 2.2 Data processing and integration

Microarray expression data for GSE32225 (Sia et al., 2013), GSE26566 (Andersen et al., 2012), and GSE132305 (Montal et al., 2020) (**Table 2**) were downloaded and processed using *GEOquery* version 2.54.1 (Sean and Meltzer, 2007). Gene annotations were updated using the R packages *AnnotationDbi* (v1.68.0) and *org.Hs.eg.db* (v3.20.0). Heatmaps were generated using *ComplexHeatmap* package version 2.2.0. Published gene expression signatures were used to reclassify tumors in the Sia *et al*. (Sia et al., 2013)(iCCA) and Andersen *et al*. (Andersen et al., 2012) (CCA) datasets according to their reported molecular and prognostic subclasses. Briefly, in the Sia *et al*. cohort (Sia et al., 2013), the proliferation and inflammation subclasses were provided in the original publication. On the other hand, survival-and recurrence-related subclasses were reassigned based on the authors’ outcome-specific signatures as described in the original study, to discriminate patients according to recurrence and survival independently of molecular subtype **(Supplementary Table 5)**. In this approach, the relative enrichment of a predefined set of genes within each sample is quantified using single-sample gene set enrichment analysis (ssGSEA), which computes an enrichment score reflecting the degree to which the genes in a given signature are coordinately up-or downregulated. Samples are then assigned to the corresponding subclass based on their relative ssGSEA scores, reproducing the stratifications described in the original studies. The resulting subclass groups were subsequently used to evaluate the distribution and expression patterns of genes and gene sets within each dataset. For the Andersen *et al*. cohort (Andersen et al., 2012), which lacked explicit subclass annotations, tumors were stratified into subclasses 1 and 2 using the published gene signature **(Supplementary Table 6**) using the same ssGSEA approach previously mentioned to stratify patients according to transcriptomic signatures. Since the Andersen et al. cohort includes both iCCA and eCCA cases and no significant prognostic differences were reported between them, we further refined sample classification using liver-specific ssGSEA scores with a gene signature defined by Hsiao et al. (Hsiao et al., 2001) **(Supplementary Table 6**). Samples with reduced hepatocyte-related enrichment were considered likely to represent eCCA, whereas higher enrichment indicated iCCA. This reclassification allowed us to account for differences in hepatic lineage contribution while maintaining the integrity of downstream analyses.

**Table 2.**
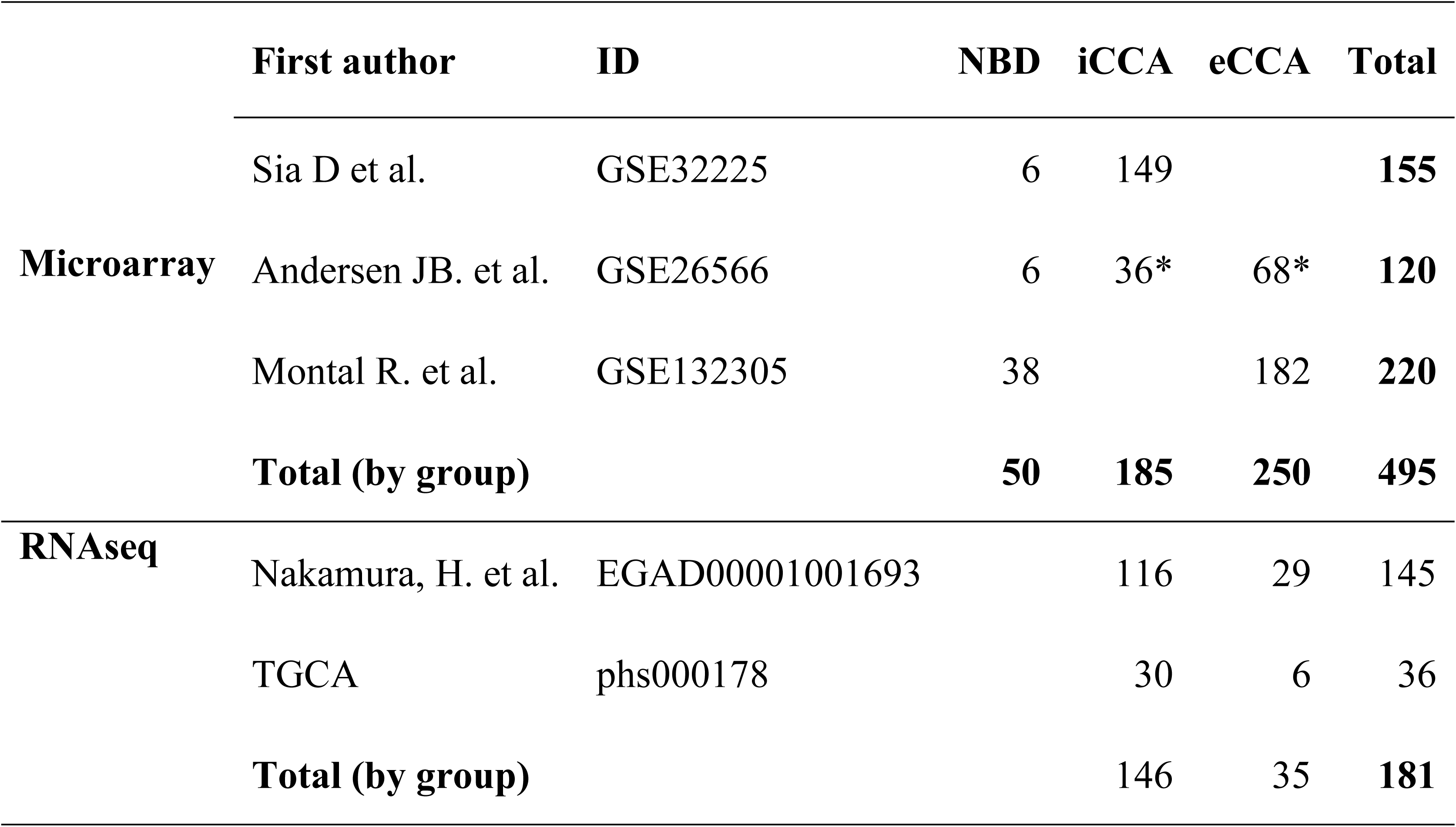
Studies included for the analysis of human liver transcriptomic data from publicly available datasets.

|  | **First author** | **ID** | **NBD** | **iCCA** | **eCCA** | **Total** |
| --- | --- | --- | --- | --- | --- | --- |
| **Microarray** | Sia D et al. | GSE32225 | 6 | 149 |  | **155** |
|  | Andersen JB. et al. | GSE26566 | 6 | 36* | 68* | **120** |
|  | Montal R. et al. | GSE132305 | 38 |  | 182 | **220** |
|  | **Total (by group)** |  | **50** | **185** | **250** | **495** |
| **RNAseq** | Nakamura, H. et al. | EGAD00001001693 |  | 116 | 29 | 145 |
|  | TGCA | phs000178 |  | 30 | 6 | 36 |
|  | **Total (by group)** |  |  | **146** | **35** | **181** |

Regarding human RNA-seq datasets (**Table 2**), gene expression data and patient clinical information were retrieved from their respective repositories. TCGA-CHOL (phs000178) gene expression data were retrieved as STAR-counts aligned to hg38 using the *TCGAbiolinks* R package (version 2.34) in R (version 4.4.2). Corresponding clinical data were obtained from the TCGA-CDR Pan-Cancer Clinical Data Resource. Raw sequencing data for EGAD00001001693 were obtained from the European Genome-phenome Archive (EGA, https://ega-archive.org/) following approval by the corresponding Data Access Committee and downloaded via the EGA Data Download Client with standard encryption and authorization procedures. EGA gene expression data were retrieved as STAR-counts aligned to hg38. The pipeline included quality control and preprocessing of the transcriptomic data as previously described (Herranz et al., 2023). Briefly, adapter sequences and low-quality reads were removed using *TrimGalore* version 0.6.0 with *Cutadapt* version 1.18 (Martin, 2011; Kechin et al., 2017). Then, the splice-aware aligner STAR version 020201 was used to align the clean reads to the reference genome with genome version hg38 as reference (Dobin et al., 2013). Aligned reads overlapping gene exons were counted using *HTseq* version 0.11.0 (Anders et al., 2015). For downstream analysis, *EdgeR* version 3.28.1 (Robinson et al., 2009) in R software version 3.6.3 (hereafter referred to as R) was used to normalize raw read counts using the trimmed mean of M-values (TMM) method (Robinson et al., 2009), which effectively addressed differences in library size and composition biases. Following the proper normalization of all samples, *ComBat* (Johnson et al., 2007) from the sva R package version 3.44.0 (Leek et al., 2012) was employed to integrate and harmonize high-dimensional biological data from the two sources, thereby reducing batch effects as previously described (Herranz et al., 2023). The integrated dataset, with diminished batch effects, was used for downstream analyses with enhanced accuracy and reliability. Furthermore, a positive-value matrix was generated by adding the minimum expression value for genes that were negative in certain samples. When genes were not detected in any of the RNA-seq datasets, they were excluded from the integrated dataset.

### 2.3 Gene expression analyses

Aligned reads are then quantified at the gene level using HTseq version 0.11.0. *EdgeR* version 3.28.1 for R software version 3.6.3 (hereafter referred to as R) requires raw read counts as input and performs normalization using the trimmed mean of M-values (TMM) method. TMM normalization accounts for library size differences and composition biases, ensuring accurate comparisons between samples. To assess transcriptomic differences among EpiGs and MGs, *edgeR* employed a generalized linear model (GLM) on normalized count data, utilizing the negative binomial distribution to account for both biological and technical variability. The identification and exclusion of low-expression genes were performed through the *filterByExpr* function in *edgeR* with default settings. Dispersion parameters were estimated using the empirical Bayes method, and statistical tests were conducted via the likelihood ratio test or the quasi-likelihood F-test.

Tumor microenvironment (TME) was evaluated using the MCP-counter method, which quantifies the relative abundance of immune and stromal cell populations directly from bulk transcriptomic data (Becht et al., 2016; Momeni et al., 2023). MCP-counter computes cell-type-specific enrichment scores based on predefined gene signatures, enabling robust estimation of T cells, CD8 T cells, cytotoxic lymphocytes, B lineage cells, NK cells, monocytic lineage, myeloid dendritic cells, neutrophils, fibroblasts, and endothelial cells. From these scores, we aggregated three functional axes reflecting key TME components: adaptive immunity (T cells, CD8 T cells, Cytotoxic lymphocytes, B lineage), innate immunity (NK cells, Monocytic lineage, Myeloid dendritic cells, Neutrophils), and stromal cells (Fibroblasts, Endothelial cells). Therefore, each CCA patient sample was represented by a three-dimensional immune-stromal vector. These vectors were used as input for unsupervised clustering, enabling patient stratification based on their global immune-stromal composition. This clustering led us to identify four subtypes of patients, which we named Immunogenic, Myeloid, Immune Desert, and Mesenchymal, based on their three-dimensional immune-stromal expression patterns, as reported in previous studies using this method applied to CCA (Job et al., 2020; Ruffolo et al., 2022; Martin-Serrano et al., 2023; Zhang et al., 2023; Zhong et al., 2024).

Gene set enrichment analysis (GSEA) was performed with *clusterProfiler* version 4.10.1 using all differentially expressed genes (raw p<0.05) without applying effect-size thresholds. Enrichment was assessed across the GO Biological Process, KEGG, and MSigDB Hallmark databases under default settings, and significance was evaluated using Benjamini-Hochberg correction. Only pathways with an adjusted p-value<0.05 were considered for interpretation.

### 2.4 Experimental models

#### 2.4.1 Cell culture models

Cell viability upon gene disruption was evaluated using publicly available CRISPR/Cas9 screens from the DepMap portal (https://depmap.org/portal/). Fitness scores, which reflect the effect of individual gene knockout on cell survival and proliferation, were obtained for 28 human CCA cell lines. Genes with a significant impact on fitness were identified, and those with a fitness score threshold of ±0.25, corresponding to a ≥25% change in viability, were used to identify genes with a substantial effect on cell survival.

The HuCCT-1 CCA cell line, of intrahepatic origin (Zach et al., 2015), was cultured in DMEM-F12 culture medium supplemented with 10% fetal bovine serum, glutamine and antibiotics, all from Gibco-Thermo Fisher (Waltham, MA, USA) as previously described (Colyn et al., 2021). Where indicated, cells were grown under hypoxic conditions (1% O_2_) using a Whitley H35 Hypoxystation (Don Whitley Scientific Ltd., Bingley, UK).

#### 2.4.2 Mouse models

Cholangiocarcinogenesis was induced in 5-week-old C57/BL6 male mice by hydrodynamic tail vein injection (HTVi) of plasmids encoding the Sleeping Beauty transposase (SB, 0.8 µg/mouse) and the oncogenic factors Notch Intracellular Domain 1 (NICD1, 10 µg per mouse) or mutant TAZ (TAZS89A, 10 µg/mouse), in combination with myr-AKT (10 µg/mouse) (GenScript, Piscataway, New Jersey, USA), which are well known to induce CCA development (Banales et al., 2020; Cigliano et al., 2022; Ma et al., 2024).

### 2.5 RNA sequencing

Total RNA from cells and tissues was extracted using the automated Maxwell system (Promega). RNA quantity and quality were assessed with the Qubit HS RNA Assay Kit (Thermo Fisher Scientific) and 4200 Tapestation with High Sensitivity RNA ScreenTape (Agilent Technologies). All samples were high-quality (RIN>8). Libraries were prepared from 100 ng of total RNA using the Illumina Stranded mRNA Prep Ligation kit, following the manufacturer’s protocol. Briefly, poly(A) RNA was captured with oligo(dT) magnetic beads, fragmented, and reverse transcribed into first-strand cDNA using random primers. Second-strand synthesis incorporated dUTP to maintain strand specificity. cDNA fragments were purified with AMPure XP beads (Beckman Coulter), adenylated, ligated to uniquely indexed adapters, purified again, and PCR-amplified. Library quality and quantity were verified with Qubit dsDNA HS Assay Kit and 4200 Tapestation with High Sensitivity D1000 ScreenTape. Libraries were sequenced on a NextSeq2000 instrument (Illumina). Adapter sequences and low-quality reads were removed using TrimGalore (v0.6.0) with Cutadapt (v1.18), and reads were aligned to mm10 as a reference genome using STAR (v2.7.9a).

### 2.6 Metabolomic profiling

Metabolomic profiling was performed by Rubió Metabolomics, S.L.U. (Bizkaia, Spain) on liver samples from control and TAZ/Akt-induced CCA in mice. Briefly, metabolites were extracted by fractionating samples according to physicochemical properties using organic solvents. Four optimized UHPLC-MS platforms were applied to achieve broad metabolome coverage(Barr et al., 2012), enabling the profiling of: fatty acyls, bile acids, steroids, and lysoglycerophospholipids; glycerolipids, glycerophospholipids, sterol lipids, and sphingolipids; amino acids; and polar metabolites. Data processing generated a list of chromatographic peak areas corresponding to the metabolites detected in each sample. For each metabolite, an approximated linear detection range was established, assuming comparable detector responses within metabolites of the same chemical class, represented by a single standard compound.

### 2.7 Proteomic analysis

Liver tissues were mechanically disrupted using a pellet pestle cordless motor in 200 μl ice-cold RIPA buffer (30 mM Tris pH 7.5, 150mM NaCl, 0.1% SDS, 0.5% sodium deoxycholate, 5mM EDTA, 1% NP-40, 1% Triton-X 100, 3.6% B-glycerophosphate, 0.5% sodium deoxycholate, 10 mM sodium fluoride and 1mM sodium orthovanadate, all purchased from Sigma) supplemented 1x protease inhibitors (Complete Mini Protease Inhibitor Cocktail, Roche). The lysates were subsequently clarified by ultracentrifugation at 100,000 ×g for 30 minutes at 4°C using a Hitachi ultracentrifuge. The resulting supernatant, corresponding to the soluble protein fraction, was carefully collected and transferred to fresh tubes for the subsequent precipitation in Methanol/Chloroform. The pellets were resuspended in 200 µL of 2.5% (w/v) SDS, 25 mM Triethylammonium bicarbonate (TEAB), 5 mM TCEP, and 10 mM chloroacetamide (CAA) supplemented with Pierce™ DNase (25 kU, 88701, ThermoFisher). Lysates were incubated at 60°C for 30 min to reduce and alkylate protein cysteine residues, followed by sonication using an ultrasonic processor UP50H (Hielscher Ultrasonics) for 1 min on ice (0.5 cycles, 100% amplitude). The protein extracts were centrifuged at 18,400 ×g for 10 min, and the supernatant was transferred to a new tube and quantified using Pierce™ 660 nm Protein Assay Reagent supplemented with Ionic Detergent Compatibility Reagent according to the manufacturer’s instructions. Automated SP3-based protein digestion was performed on the Opentrons OT-2 platform in the presence of MagReSyn® Hydroxy microparticles as described elsewhere (Ciordia et al., 2024), with the corresponding adaptations for our sample format. Proteins were digested overnight at 37°C using trypsin (1:33, enz:prot) and Lys-C (1:500, enz:prot). Eluted peptides were dried by speed vacuum and quantified by fluorimetry (Qubit). Peptide samples were labeled using the TMT-18plex Isobaric Mass Tagging Kit according to the manufacturer’s instructions, including two pooled internal standards labeled with the 133C and 133N channels. After labeling, samples were pooled, evaporated to dryness, and stored at-20°C. High-pH reversed-phase fractionation was performed using Styrene Divinylbenzene reverse phase sulfonate (SDB-RPS) StageTips (CDS Empore™, Sigma-Aldrich) as previously reported (Ciordia et al., 2024), applying stepwise elution with increasing acetonitrile (ACN) concentrations (0-60%) in 10 mM ammonium formate (pH 10.0) to obtain five fractions, which were dried and stored at-20 °C.

For nano-Liquid Chromatography coupled to Electrospray Ionization Tandem Mass Spectrometry (nanoLC-ESI-MS/MS) analysis, 500 ng of each high-pH fraction was injected into an Ultimate 3000 nano-HPLC system (Thermo Fisher Scientific) connected online to an Orbitrap Exploris™ 240 mass spectrometer as previously reported (Ciordia et al., 2024). Briefly, peptides were separated on an Easy-spray PepMap C18 analytical column. MS1 spectra were acquired at 60,000 resolution (m/z 200), followed by data-dependent selection of the top 20 precursors for higher-energy collisional dissociation fragmentation (34% NCE). MS2 spectra were acquired at 45,000 resolution with an automatic gain control (AGC) target of 50%, using a 0.7 m/z isolation window and a 45-s dynamic exclusion. Precursor ions with charge states from +2 to +5 were selected for fragmentation.

### 2.7 Statistical analyses

#### 2.7.1 Gene expression statistical analyses

Genes were considered differentially expressed if the adjusted p-value (FDR method of Benjamini and Hochberg) was lower than 0.05. For exploratory analyses within predefined lists of EpiGs and MGs, all genes with a raw p-value<0.05 were considered deregulated. Multiple comparisons of gene set categories were controlled by the False Discovery Rate (FDR) using the Benjamini and Hochberg correction (Q=5%). For comparisons between two groups, data were tested for normality and homogeneity of variances. Normally distributed data with homogeneous variances were analyzed using a two-tailed unpaired Student’s t-test, while Welch’s correction was applied when variances were unequal. For nonparametric data, the Mann-Whitney U test or the Kolmogorov-Smirnov test was used, depending on whether the variances were homogeneous. Regarding human organoids and tumoroids (Broutier et al., 2017), the number of biological replicates was limited to account for human heterogeneity (n=3 per group), and the statistical power to detect differentially expressed genes (DEGs) was low, resulting in few genes reaching conventional significance thresholds. Therefore, DEGs with *p*-values<0.25 and an absolute Log2FC>0.10 are considered here for exploratory purposes, to facilitate the identification of potential candidates for further validation. Gene expression correlation was computed using Pearson correlation methods. GraphPad Prism 9.0.2 software (GraphPad Prism, San Diego, CA, USA) was used for these statistical analyses and the corresponding boxplots. Data are presented as mean and standard error. The differentially expressed gene sets were depicted using the InteractiVenn tool (Heberle et al., 2015). Values of p<0.05 were considered statistically significant.

#### 2.7.2 Metabolites statistical analyses

In the metabolomic studies, normalization of the dataset was performed according as previously described (Martínez-Arranz et al., 2015). To reduce the dimensionality of the dataset and allow visualization of potential clustering between experimental groups, multivariate data analysis was conducted using unsupervised principal component analysis (PCA). Analyses were performed in SIMCA software (version 14.1, MSK Umetrics AB, Umea, Sweden), with the data mean-centered and unit-variance (UV)-scaled. Model quality was assessed by calculating R², which represents the proportion of variance explained by the model, and Q², which reflects predictive accuracy, both estimated using 7-fold cross-validation. In addition, univariate statistical analyses were performed for each metabolite. Data distribution was first evaluated using the Shapiro-Wilk test to assess normality. Depending on the distribution, differences between groups were determined using Student’s t-test (for normally distributed data) or the Wilcoxon-Mann-Whitney U test (for non-normal data). Fold-change values were also calculated to evaluate the magnitude of differences.

#### 2.7.2 Protein statistical analyses

Raw data files were processed using the Proteome Discoverer 2.5.0.400 software (Thermo Scientific, Bremen, Germany), and a database search was carried out using using three search engines (Mascot (v2.8.0), MsFragger (v3.1.1), and Sequest HT) against *Mus musculus* UniProtKB reviewed database (14th September 2024, 20,470 sequences) containing the most common laboratory contaminants (cRAP database with 70 sequences). Search parameters were set as follows: cysteine carbamidomethyl (+57.021464 Da) and TMTpro (+304.207 Da) on lysine and N-term as fixed modifications; methionine oxidation (+15.995 Da), N-term acetylation (+42.010 Da), and Gln→pyro-Glu (-17.026 Da) as variable modifications. Precursor mass tolerances were set at 10 ppm and the fragment mass tolerance at 0.02 Da and trypsin/P was selected as a protease with a maximum of 2 missed cleavage sites. False discovery rate (FDR) was calculated using the processing node Percolator (maximum delta Cn 0.05; decoy database search target) and the validation of proteins, peptides, and peptide spectral matches (PSMs) peptides with an FDR≤1 %. The quantitation was also performed in Proteome Discoverer using the “Reporter Ions Quantifier” feature in the quantification workflow using the following parameters: unique+razor peptides were used for quantitation, co-isolation threshold was set at 50%, signal to noise of reporter ions was 10, and the normalization and scaling were performed considering the total peptide amount and the control (IS) average, respectively. The protein ratio was calculated considering the protein abundance and the hypothesis test was based on a “*t-test (background-based)*”. Protein groups (master proteins) with an FDR lower than 1% and with abundance values in both IS were considered for quantitation. To identify the proteins that were differentially expressed in each comparison, an adjusted p-value threshold of ≤0.05 was applied using the Benjamini-Hochberg post hoc adjustment. Volcano plot and Principal Component Analysis (PCA) were performed in Proteome Discover considering the differentially expressed proteins in each comparison.

## 3 Results

### 3.1 Transcriptional dysregulation of epigenetic genes in human CCA

Epigenetic modifiers (EpiG) comprising 11 families and a total of 257 genes were selected for the analyses (**Supplementary Table 1**). These are considered the most widely described genes that belong to three different categories of epigenetic writers: DNA methyltransferases (DNMTs), protein arginine-methyltransferases (PRMTs), protein lysine-methyltransferases (KMTs), histone acetyl-transferases (HATs); epigenetic erasers: DNA demethylases (TETs), histone-lysine demethylases (HDMs), histone deacetylases (HDACs), and histone deiminases (HDIs); and epigenetic readers: DNA methyl-binding proteins (MBPs), histone methyl readers (HMRs), and histone acetyl readers (HARs) (**Table 1**). Further details on their functions, targets, and at least one relevant literature reference have been recently reported (Herranz et al., 2023; Castelló-Uribe et al., 2025), and are available in **Supplementary Table 1**. We analyzed three publicly available microarray-based transcriptomic datasets, including samples of non-neoplastic bile duct epithelia (NBD) along with iCCA and eCCA (pCCA and dCCA) tumoral tissues (**Table 2**). As observed in the heatmaps shown in **Figures 1, 2,** and **3**, marked changes in the expression of numerous specific EpiGs, mostly upregulation, were detected between CCA tissues (both iCCA and eCCA) and NBDs. The expression of selected EpiGs that were significantly changed in tumors is shown in **Supplementary Figure 1**. We validated the overexpression of genes such as *DNMT1*, *EZH2*, *UHRF1*, *SUZ12*, *BRD4*, *HDAC1* and *HDAC3* (**Supplementary Figure 1A**) as previously reported by us and others (Yin et al., 2017; Wasenang et al., 2019; Colyn et al., 2021; Hu et al., 2022; Xu et al., 2022; Zhang et al., 2022; Wu et al., 2023; Chen et al., 2024; Xiong et al., 2024). We also observed the upregulation of additional EpiGs that may be relevant to CCA growth and development, including readers such as *CBX3*, *PHF20L1, SMARCA4*; writers like *SMYD3*, and erasers as *KDM5C* (**Supplementary Figure 1B**). However, some EpiGs, such as the readers *CHD5* and *SMARCA2*, were consistently downregulated in CCA (**Supplementary Figure 1C**).

**Figure 1.**
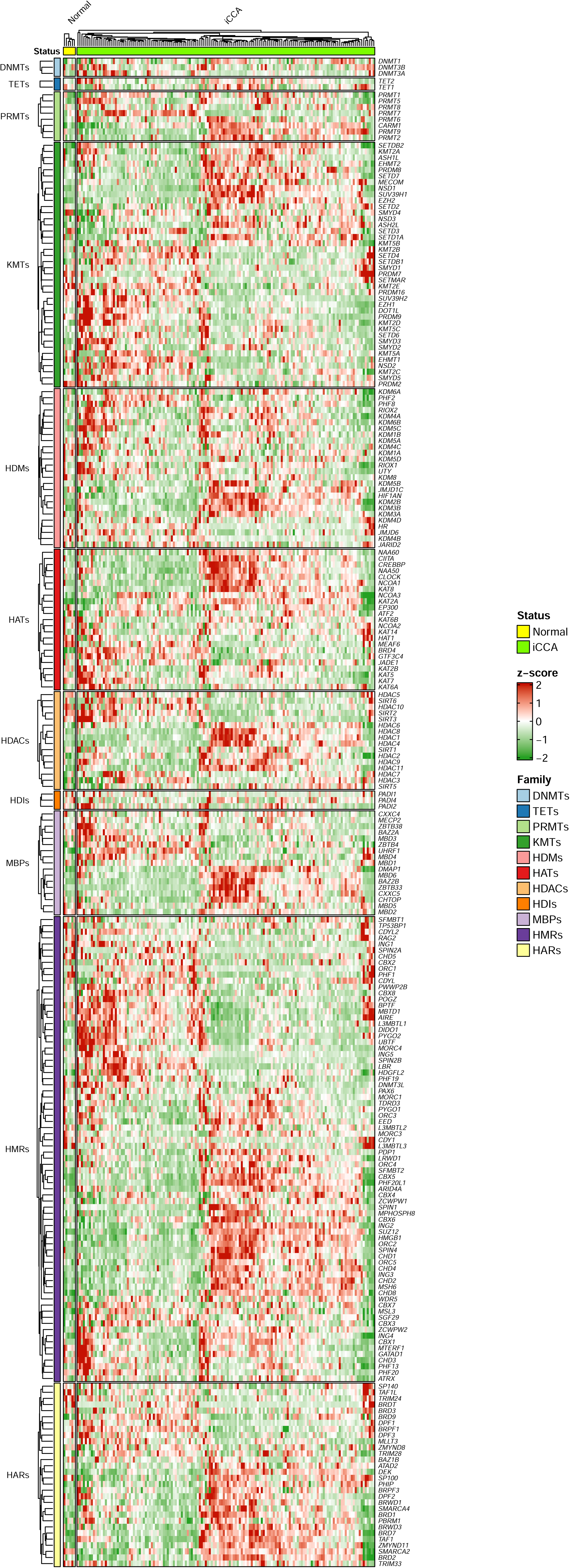
EpiGs dysregulation in human iCCA. Heatmap of the expression of epigenetic genes (EpiGs) grouped in families in non-neoplastic bile duct epithelia (NBD) and intrahepatic cholangiocarcinoma (iCCA) in the GSE32225 dataset.

**Figure 2.**
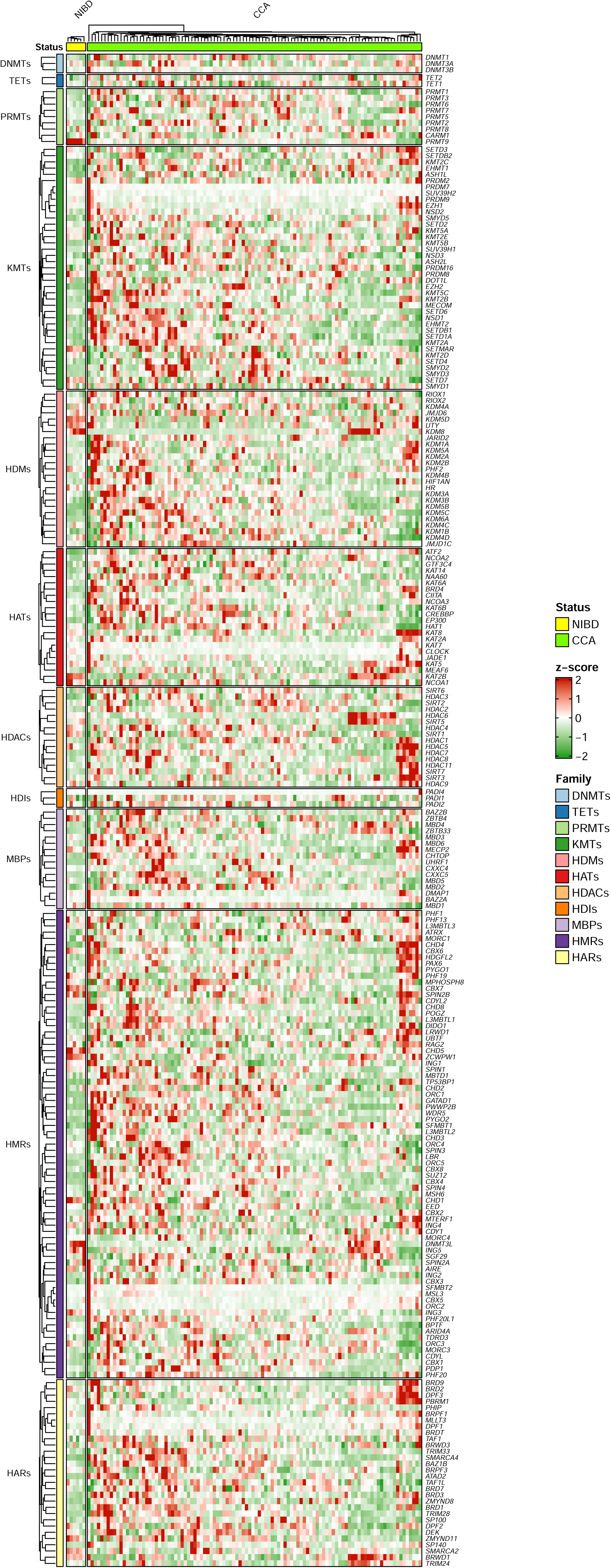
EpiGs dysregulation in human CCA. Heatmap of the expression of epigenetic genes grouped in families in non-neoplastic bile duct epithelia (NBD) and cholangiocarcinoma (CCA) in the GSE26566 dataset.

**Figure 3.**
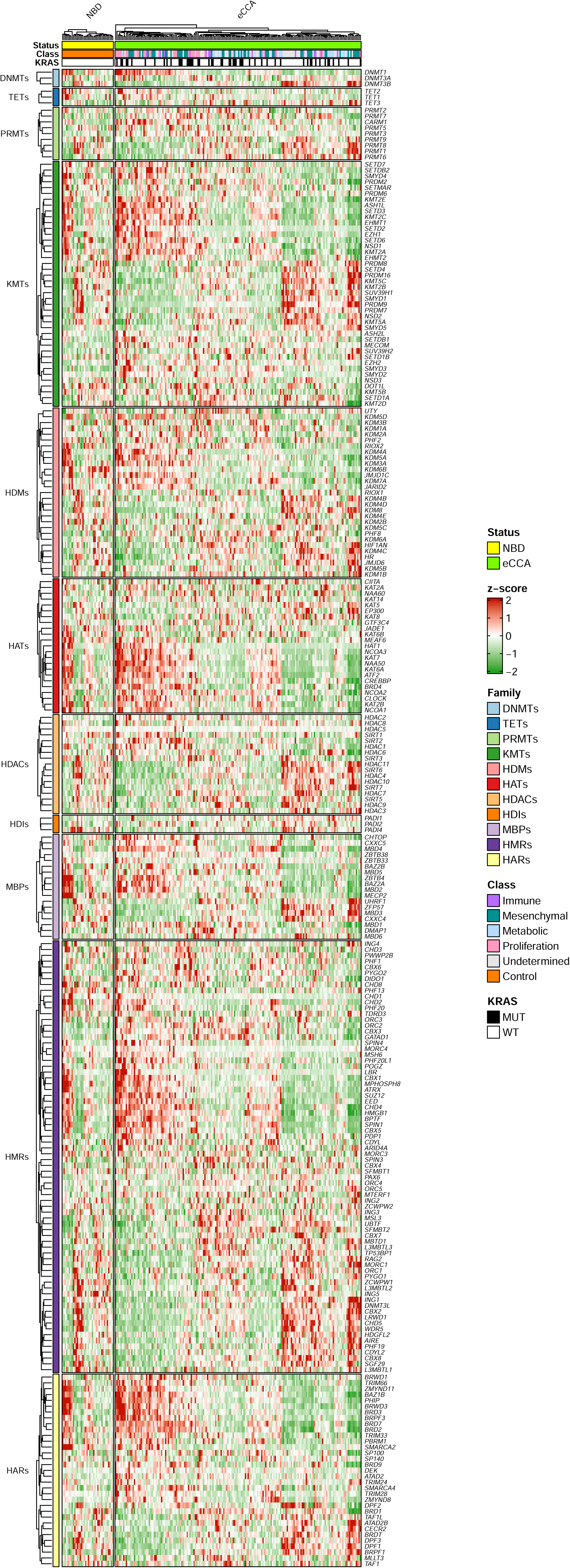
EpiGs dysregulation in human eCCA. Heatmap of the expression of epigenetic genes grouped in families in non-neoplastic bile duct epithelia (NBD) and extrahepatic cholangiocarcinoma (eCCA) in the GSE132305 dataset.

To explore potential epigenetic contributions to the transcriptional alterations observed in CCA, we analyzed DEGs using GO, KEGG, and Hallmark gene sets in GSEA, highlighting pathways associated to epigenetically related mechanisms (**Supplementary Table 7**). Thus, we selected enriched terms (over-or underrepresented) with statistically significant Normalized Enrichment Score (NES) (adjusted *p*<0.05) that included EpiGs among the DEGs, noting that nearly all such gene sets were found among the overrepresented pathways. This indicates that epigenetic mechanisms are predominantly associated with upregulated processes in CCA. Gene sets containing ten or more EpiGs and consistently overrepresented in at least two datasets were mainly related to DNA repair, epigenetic regulation of gene expression, and chromatin organization, including the biological processes “regulation of DNA repair” and “heterochromatin formation”, the molecular function “modification-dependent protein binding”, and “chromosomal-related compartments”. In addition to the broader pathways containing ≥10 EpiGs—representing general epigenetic and chromatin-related functions commonly altered across the CCA datasets—several specific terms with at least three EpiGs highlighted processes, such as “cell cycle regulation”, “mitotic progression”, “nucleosome organization”, “chromatin remodeling”, and “transcriptional elongation”, further support the involvement of epigenetic regulators in the transcriptional reprogramming of CCA. A subset of pathways was consistently overrepresented across all three datasets, reflecting recurrent epigenetic-related transcriptional programs (**Supplementary Figure 2A**). These included nucleosome-and chromatin-associated terms (“nucleosome organization”, “protein-DNA complex”, “protein localization to chromatin” and “chromosome, chromosomal and centromeric regions”), “ribonucleoprotein complex biogenesis”, “viral processes”, and “key proliferative signatures” (G2M checkpoint, E2F targets, MYC targets V1). The consistent upregulation of these pathways underscores the central role of epigenetic mechanisms in shaping the transcriptional landscape of CCA.

### 3.2 Human CCA metabolic transcriptional reprogramming

We next evaluated the expression of 96 MGs encoding metabolic enzymes functionally linked to epigenetic regulation (**Table 1**), given their role in generating essential cofactors that modulate the activity of epigenetic writers, erasers, and readers. These enzymes participate in the synthesis and metabolism of cofactors involved in the activity and regulation of most epigenetic reactions (**Supplementary Table 2**) (Li et al., 2018; Reina-Campos et al., 2019; Esteve-Puig et al., 2020; Boon, 2021). **Figure 4** summarizes the expression patterns of these metabolic genes across datasets, highlighting those significantly dysregulated in CCA vs. NBD. Focusing on genes significantly up-or downregulated in at least two of the three studies, and not oppositely regulated in the third, we found six upregulated MGs in CCA: within the OCM pathway *ALDH1L2*, *ATIC*, *GART*, and *TYMS*, and in the TCA-cycle *IDH2*, and *SDHC*; and 16 downregulated MGs: in the OCM pathway *ALDH1L1*, *AMT*, *BHMT*, *FTCD*, *GNMT*, *MAT1A*, *NNMT*, *PSAT1*, and *SHMT1*, in the TCA-cycle *MDH1B*, *PCK1*, *PDHA2*, and *PDK4*, and finally in the ACS pathway *ACSM2A*, *ACSM2B*, and *ACSM6*.

**Figure 4.**
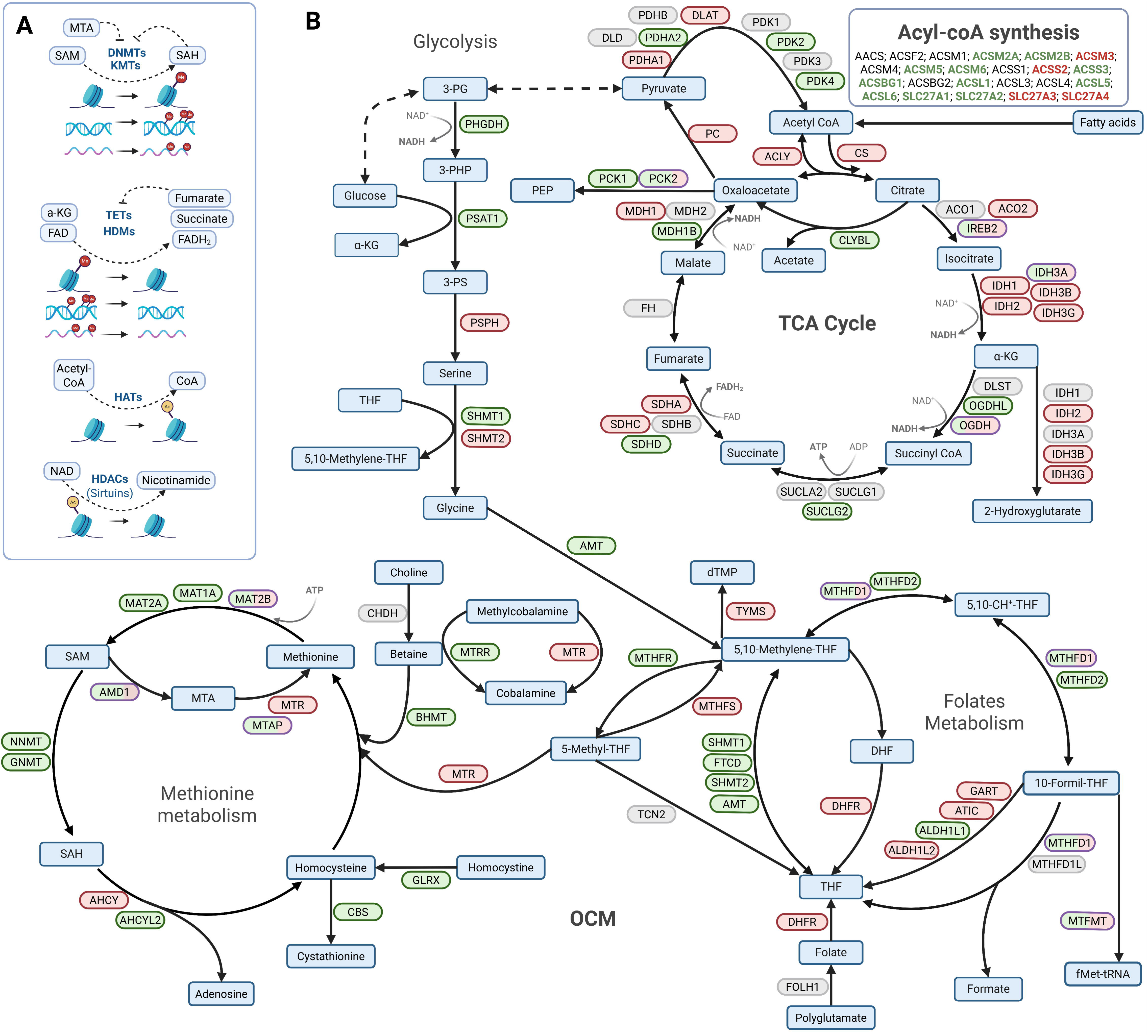
Metabolic-epigenetic cross-talk with pathway-level gene expression alterations in human CCA. **(A)** Cross-talk between epigenetic factors and metabolites acting as substrates or inhibitors of these reactions. **(B)** Major metabolic pathways involved in the synthesis and conversion of substrates and cofactors required for epigenetic enzymatic activity. Gene expression fold changes (Log2FC) are shown as green (downregulated) or red (upregulated) in cholangiocarcinoma (CCA) samples compared with normal bile ducts (NBDs) in datasets GSE32225, GSE26566, and GSE132305. The colored fraction of each symbol represents the proportion of datasets in which a given gene is significantly deregulated (raw p<0.05). Created in https://BioRender.com. Abbreviations can be found in the Supplementary Tables 1 and 2.

Furthermore, the expression of more than 1,000 genes associated with 92 metabolic pathways, previously reported to display distinct patterns between tumor and normal tissues (Rosario et al., 2018), was examined as an extended analysis of metabolic reprogramming (**Supplementary Table 3**). CCA datasets uncovered a striking and consistent downregulation of multiple metabolic pathways, spanning amino acid, lipid, carbohydrate, and xenobiotic metabolism (**Supplementary Table 7**). Pathways related to oxidative phosphorylation, glycolysis, ribosome biogenesis, aminoacyl-tRNA biosynthesis, and proteasome function were markedly enriched (**Supplementary Table 7**), indicating an increased demand for energy and macromolecule synthesis. These changes point to a metabolic rewiring in malignant cholangiocytes characterized by enhanced mitochondrial activity, increased translational capacity, and proteostatic control, consistent with a coordinated adaptation linking metabolic flux to epigenetic regulation. A subset of metabolism-related pathways was consistently dysregulated across datasets (**Supplementary Figure 2A**). Underrepresented terms included processes linked to ion homeostasis, metabolite transport, and redox balance (“cellular response to copper ion”, “detoxification of inorganic compounds”, “organic acid and anion transport”, “one-carbon compound transport”), while overrepresented pathways reflected enhanced energy production and biosynthesis (glycolysis, mitochondrial gene expression and translation, lysosome, proteasome, PI3K-AKT-mTOR signaling). Overall, this metabolic profile reflects a broad suppression of normal biliary functions and aligns with the dedifferentiated state of CCA cells, which favor proliferative and survival programs over canonical metabolic pathways.

Analysis of 189 RLEs expression **(Supplementary Table 4**) showed consistent enzyme deregulation (**Supplementary Table 8**) with 13 upregulated and 25 downregulated genes encoding for RLEs in at least two of the three studies, and not oppositely regulated in the third. We further analyzed DEGs encoding RLEs whose up-or downregulation (at least in two datasets) was consistent with the metabolic pathways identified as over-or underrepresented in CCA. Upregulated RLEs were mainly involved in nucleotide synthesis and redox metabolism, including *G6PD* and *TKT* (pentose phosphate pathway), *RRM2*, *TYMS*, and *TYMP* (pyrimidine and purine metabolism), and *PKM* (pyruvate metabolism), indicating enhanced anabolic and antioxidant activity. In contrast, downregulated RLEs such as *ACACB* and *ACADL* (fatty acid metabolism), *CPS1*, *CTH*, *PCK1*, and *FBP1* (amino acid and gluconeogenic pathways), and *LIPC*, *LIPG*, and *LPL* (glycerolipid metabolism) reflected suppression of catabolic and hepatobiliary-specific metabolic processes. These findings highlight a coordinated shift toward biosynthetic and redox-supporting pathways alongside the loss of oxidative and detoxifying functions in CCA, which is consistent with the suppression of lipid, amino acid, and xenobiotic metabolism observed at the pathway level.

### 3.3 Transcriptomic patterns distinguishing iCCA and eCCA

In line with the distinct cellular and genetic characteristics of iCCAs and eCCAs (Brindley et al., 2021; Ilyas et al., 2023), we identified 440 DEGs with opposite expression patterns between GSE32225 (Sia et al., 2013)(iCCA vs NBD) and GSE132305 (Montal et al., 2020)(eCCA vs NBD). Among these, a subset of EpiGs displayed inverse expression changes in both tumor types (**Table 3** and **Supplementary Table 8**). Specifically, among the genes upregulated in iCCA (Sia et al., 2013) but downregulated in eCCA (Montal et al., 2020) were the readers *L3MBTL3* and *MTERF1*. Conversely, genes downregulated in iCCA but upregulated in eCCA included three readers, *PHF13*, *CHD2*, and *CHD8*, as well as three erasers, *KDM2B*, *SIRT5*, and *HDAC4*. These patterns suggest that opposing epigenetic regulation may contribute to the molecular distinctions between iCCA and eCCA. Among the MGs encoding metabolic enzymes functionally linked to epigenetic regulation, two (*AMD1* and *MTAP)* were upregulated in iCCA (Sia et al., 2013) but downregulated in eCCA (Montal et al., 2020), and six (*MAT2B*, *MTFMT*, *MTHFD1*, *IREB2*, *OGDH*, *PCK2*) showed the opposite trend, being downregulated in iCCA (Sia et al., 2013) but upregulated in eCCA (Montal et al., 2020) (**Supplementary Table 8**). Regarding the RLEs analyzed (**Supplementary Table 8**), four genes encoding these RLEs exhibited a marked opposite expression between the iCCA (Sia et al., 2013) and eCCA (Montal et al., 2020) datasets (*APRT*, *GPAT4, OGDH*, and *PPAT*). Comparison of the GSEA results from the GSE32225 (Sia et al., 2013)(iCCA vs NBD) and GSE132305 cohorts (Montal et al., 2020)(eCCA vs NBD) identified 38 pathways displaying opposite enrichment directions (adjusted p<0.05) (**Supplementary Figure 2B**). Of these, 30 pathways were underrepresented in iCCA (Sia et al., 2013) but overrepresented in eCCA (Montal et al., 2020), mainly encompassing lipid and lipoprotein metabolism, detoxification and xenobiotic processing, coagulation, and extracellular matrix-related functions. In contrast, eight pathways were overrepresented in iCCA (Sia et al., 2013) but underrepresented in eCCA (Montal et al., 2020), including stress-and signaling-related processes (TNFα signaling via NF-κB, UV response, apoptosis), nuclear organization, and protein interaction networks. Together, these findings reveal a clear divergence in pathway enrichment between both datasets, exhibiting relative enrichment of stress response and signaling pathways and suppression of lipid, detoxification, and extracellular matrix-associated programs in iCCA (Sia et al., 2013) compared with eCCA (Montal et al., 2020).

**Table 3.** Epigenetic genes with opposite expression in iCAA and eCCA.

| EpiG |  |  | Sia *et al.* |  |  | Montal *et al.* |  |  |
| --- | --- | --- | --- | --- | --- | --- | --- | --- |
| Symbol | Family | Class | LFC | *p*-value | FDR | LFC | *p*-value | FDR |
| *L3MBTL3* | HMRs | Reader | -0.56 | 7.50E-04 | 6.51E-03 | 0.05 | 1.44E-02 | 7.64E-02 |
| *MTERF1* | HMRs | Reader | -0.20 | 3.39E-02 | 1.02E-01 | 0.05 | 3.90E-02 | 1.49E-01 |
| *KDM2B* | HDMs | Eraser | 0.27 | 3.88E-02 | 1.12E-01 | -0.07 | 2.45E-03 | 2.19E-02 |
| *PHF13* | HMRs | Reader | 0.33 | 2.07E-02 | 7.17E-02 | -0.14 | 6.73E-05 | 1.39E-03 |
| *SIRT5* | HDACs | Eraser | 0.50 | 1.74E-04 | 2.26E-03 | -0.06 | 2.89E-02 | 1.22E-01 |
| *HDAC4* | HDACs | Eraser | 0.69 | 5.38E-03 | 2.72E-02 | -0.12 | 1.59E-04 | 2.72E-03 |
| *CHD2* | HMRs | Reader | 0.89 | 4.78E-04 | 4.69E-03 | -0.26 | 6.52E-03 | 4.42E-02 |
| *CHD8* | HMRs | Reader | 1.14 | 1.72E-08 | 2.11E-06 | -0.09 | 2.97E-02 | 1.24E-01 |

Since the cohort published by Andersen *et al*. (Andersen et al., 2012) comprises both iCCA and eCCA cases and no significant prognostic differences were reported, we reclassified these samples according to their ssGSEA scores using the liver-specific signature defined by Hsiao *et al*. (Hsiao et al., 2001).

Assuming a reduced hepatic signature, eCCA samples were expected to display lower hepatocyte-related enrichment scores. Thus, we explored whether the transcriptional differences observed between the GSE32225 (Sia et al., 2013) (iCCA vs. NBD) and GSE132305 (Montal et al., 2020) (eCCA vs. NBD) datasets could also be detected within the Andersen cohort by including in our analyses the reassigned Hsiao_Low vs. NBD and Hsiao_High vs. NBD samples from GSE26566 (Andersen et al., 2012) (**Supplementary Table 6**). We then examined the DEGs, focusing on overall expression patterns, EpiGs, MGs, and genes encoding for RLEs. Looking at the overlapping genes between datasets, we identified a subset of 230 genes consistently upregulated and 178 downregulated genes across all comparisons, reflecting shared transcriptional programs in CCA (**Supplementary Figure 2 C-D**). Notably, several of these concordant genes corresponded to epigenetic regulators (*HDAC3*, *PHF20L1*, *CBX3*, *EZH2,* and *SMARCA4* are upregulated, while *CHD5* is downregulated in all CCA datasets), suggesting a role of these EpiGs regardless of the CCA anatomic location. Regarding the MGs coding for metabolic enzymes functionally linked to epigenetic regulation, none was downregulated across all datasets, but *GART* and *TYMS* were upregulated in all of them. Downregulated RLEs across datasets were *ADH1B* and *HDC*, while upregulated RLEs were *MAN2B1*, *PGAP6*, *PKM*, *RRM2*, and *TYMS*. Given the potential bias introduced by the peritumoral origin of the NBD samples in Montal et al. (Montal et al., 2020), this cohort (GSE132305) was excluded from subsequent analyses of aggressiveness.

### 3.4 Epigenetic and metabolic genes’ expression associate with CCA aggressiveness

According to transcriptomic patterns, Sia *et al*. detected two distinct classes of iCCAs, the proliferation and inflammation classes, with patients in the proliferation class showing poorly differentiated and more aggressive tumors (Sia et al., 2013). Similarly, by hierarchical cluster analysis, Andersen and collaborators found two tumor subclasses, 1 and 2, with subclass 2 being associated with increased cellular proliferation and worse survival (Andersen et al., 2012). Since subclass information was not explicitly annotated in the latter, we inferred the groups by applying the signature described in their study, which allowed us to reassign samples into prognostic subclasses 1 and 2 (**Supplementary Table 6**). We also reclassified the iCCA patients of GSE32225 (Sia et al., 2013) according to the survival and recurrence signatures described in this study, which allowed us to reassign samples into survival (poor vs. good prognosis), and recurrence (poor vs. good outcome) (**Supplementary Table 5**).

Differential expression analysis showed extensive remodeling of EpiGs across the main molecular subclasses of CCA. In the GSE32225 (Sia et al., 2013)(iCCA) cohort, 185, 171, and 150 EpiGs were differentially expressed (raw p<0.05) in the proliferation-inflammation, survival (poor vs. good prognosis), and recurrence (poor vs. good outcome) comparisons, respectively. Among these, 109/76, 107/64, and 89/61 genes were up-and downregulated in each comparison. When comparing the Andersen *et al*. (Andersen et al., 2012) subclasses 2 (worse prognosis) and 1 (better prognosis), we identified 72 DEGs (35 downregulated and 37 upregulated). Several EpiGs showed consistent dysregulation in the worst versus best prognosis contexts in the molecular subclasses, survival, and recurrence comparisons: downregulated genes included *BPTF, DNMT3L, ING5, LBR, MEAF6, MORC4, SETMAR,* and *SPIN2A* (**Figure 5A**), whereas *CHD4, CHD8, CIITA, HDAC1, HDAC2, MECOM, MPHOSPH8, PRDM8, PRMT1, SETDB2, SP100, SPIN4,* and *TDRD3* were recurrently upregulated (**Figure 5B**). Supervised DAPC identified a small subset of EpiGs that best discriminated between groups (loadings >2% of between-group variance) (**Figure 5C**). We also examined dysregulation of these discriminant genes by comparing Log2FC values between the poor-and favorable-prognosis groups across survival, recurrence, and molecular subclass analyses. In GSE32225 (Sia et al., 2013) proliferation vs. inflammation subclasses, the discriminant and differentially expressed EpiGs were: *HDAC2, ING2*, and *SETDB2* as upregulated, and *ING1* as downregulated. In GSE32225 (Sia et al., 2013) survival analysis, *HDAC2* was upregulated, and *KMT2E*, *POGZ*, and *PWWP2B* were downregulated. In the GSE32225 (Sia et al., 2013) recurrence comparison, *HDAC2, LRWD1*, and *PRDM8* were upregulated, while *POGZ* and *SETDB1* were downregulated. In the GSE26566 (Andersen et al., 2012) classification into subclasses 2 (worse prognosis) and 1 (better prognosis), 11 EpiGs showed DAPC loadings >2% and differential expression. Taking into consideration the discriminant and differentially expressed EpiGs between subclasses 1 and 2, we found that *HDAC1*, *MECOM*, *PRDM8*, *PRMT5, PRMT7*, and *SIRT7* were upregulated, and *CXXC5*, *LBR, MEAF6*, *SMYD2*, and *SPIN3* were downregulated. Collectively, the eraser *HDAC2* consistently emerged as a discriminant gene upregulated in the most aggressive subclasses, while the reader *POGZ* showed the opposite pattern. The recurrent involvement of *HDAC1*, *HDAC2*, *PRDM8*, and *MECOM* among both the DAPC-selected and differentially expressed EpiGs highlights a shared signature of chromatin repression and histone modification linked to poor prognosis. The pathway enrichment analyses could clarify whether these convergent genes define specific epigenetic programs associated with tumor progression. To this end, we examined enriched gene sets (adjusted p<0.05) containing at least three significantly deregulated EpiGs in the worst-versus best-prognosis groups across the GSE32225 (Sia et al., 2013) and GSE26566 (Andersen et al., 2012) cohorts **(Supplementary Table 9)**. Several overrepresented pathways containing differentially expressed EpiGs were consistently identified, encompassing “TGF-β-related signaling” (e.g., “transforming growth factor-beta receptor and transmembrane receptor serine/threonine kinase signaling”), “myeloid cell differentiation”, and “key processes related to organelle fission and chromosomal segregation”. In contrast, no shared underrepresented sets were detected between analyses. These findings indicate that the dysregulation of EpiGs in aggressive CCA subclasses converges on programs controlling proliferative signaling, mitotic progression, and transcriptional remodeling, consistent with the establishment of an epigenetic landscape that favors tumor growth and poor clinical outcome.

**Figure 5.**
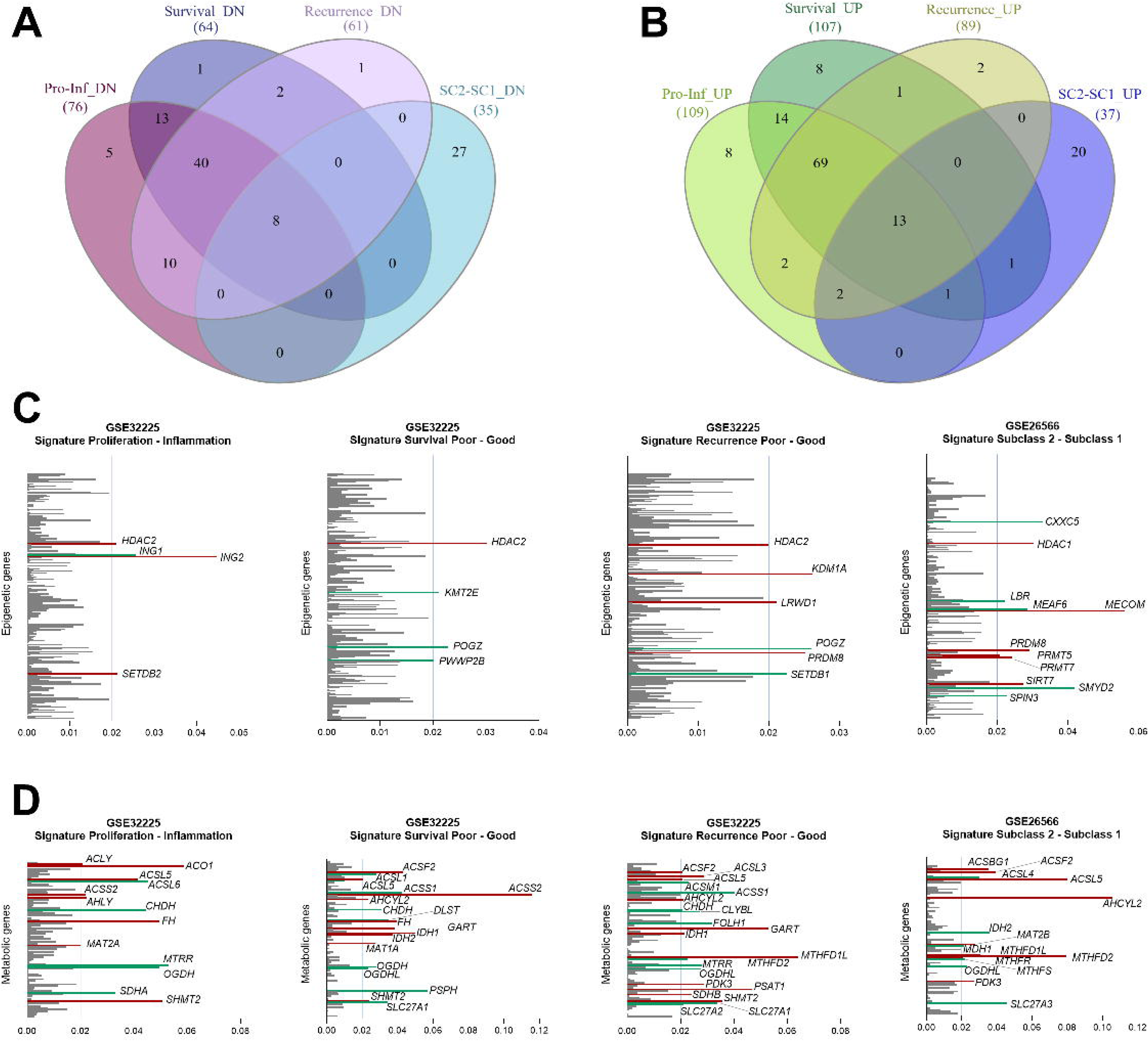
Epigenetic and metabolic expression programs associated with CCA aggressiveness transcriptomic signatures. Venn diagrams showing the overlap of identified **(A)** downregulated and **(B)** upregulated genes in patients classified into the poor-prognosis and aggressive subclasses of CCA, compared with better-prognosis subclasses, in the GSE32225 and GSE26566 datasets. **(C)** Most relevant EpiGs contributing to the stratification of patients in the prognosis and aggressive subclasses in GSE32225 and GSE26566 datasets. **(D)** Most relevant MGs contributing to the stratification of patients in the prognosis and aggressive subclasses in GSE32225 and GSE26566 datasets. In the loading plots, the colored bars indicate when genes are downregulated (green) or upregulated (red) in the corresponding dataset when comparing poor vs. good prognosis. The scale indicates each gene’s contribution to the separation between groups. Selected genes explain at least 2% of the variability between poor-and good-prognosis subtypes.

Regarding the MGs encoding metabolic enzymes linked to epigenetics, we also found consistent deregulation in the worst versus best prognosis contexts: *ACO1*, *ACSM2A*, *CBS*, *CHDH*, *FH, FOLH1*, *IDH1*, *IDH2*, *MDH1*, *OGDHL*, *PCK2*, *PDHB*, *PHGDH*, *SDHC*, *SHMT1*, *SHMT2*, *SLC27A* were downregulated, while *ACSL5*, *AHCYL2*, and *MTHFD1L* were upregulated. Supervised DAPC identified a subset of MGs with the highest discriminant loadings (>2% of between-group variance) (**Figure 5D**). Differential expression analysis between prognostic groups identified consistent metabolic alterations across classifications. In the proliferation vs. inflammation subclasses, *ACLY*, *ACO1*, *ACSL5*, *ACSS2*, *AHCY*, *FH*, *MAT2A*, and *SHMT2* were upregulated, whereas *ACSL6*, *CHDH*, *MTRR*, *OGDH*, and *SDHA* were downregulated. In the survival analysis, *ACSL5*, *AHCYL2*, *FH*, *GART*, *IDH1*, *IDH2*, *MAT1A*, and *SHMT2* were upregulated, while *ACSL1*, *ACSL6*, *ACSS1*, *CHDH, DLST*, *OGDH*, *OGDHL*, and *SLC27A1* were downregulated. In the recurrence comparison, *ACSF2*, *ACSL3*, *ACSL5*, *AHCYL2*, *GART*, *IDH1*, *MTHFD1L*, *SDHB*, and *SHMT2* were upregulated, whereas *CHDH*, *FOLH1*, *MTRR*, *OGDHL*, and *SLC27A1* were downregulated. Finally, in the Andersen subclasses, subclass 2 vs subclass 1, *ACSL5*, *AHCYL2*, *MAT2B*, *MTHFD1L*, *MTHFD2*, *MTHFR*, and *PDK3* were upregulated, while *ACSL4*, *IDH2*, *MDH1*, *MTHFS*, *OGDHL*, and *SLC27A3* were downregulated. Collectively, the MGs that were both discriminant and differentially expressed across analyses demonstrated a recurrent upregulation of one-carbon and lipid metabolic enzymes (e.g., *SHMT2*, *AHCYL2*, *ACSL5*) and the consistent downregulation of oxidative and mitochondrial components (e.g., *OGDHL*, *ACSL6*, *CHDH*) in aggressive CCA subclasses, indicating a coordinated metabolic reprogramming associated with poor prognosis.

To assess whether our gene lists (EpiGs and MGs) are associated with patients’ survival, we integrated two independent datasets: the Nakamura et al. cohort (EGA) and the TCGA cohort (details of patients and datasets are provided in **Table 2**). Using this integrated dataset, we evaluated the relationship between gene expression and survival time, with results summarized in **Table 4**. Several genes whose expression distinguished aggressive CCA subclasses also showed significant correlations with patients’ survival. Concretely, in iCCA, EpiGs such as *HDAC1*, *HDAC2*, *ING2*, *MPHOSPH8*, *PRDM8*, *SP100*, and *SPIN4*, previously linked to poor-prognosis groups, showed a negative association with survival, further supporting their role in tumor aggressiveness. Among the MGs, we found that *ACSF2*, *ACSL5*, *MTHFD1L*, *MTHFD2*, and *PDK3*, which were recurrently upregulated in aggressive subclasses, also correlated with shorter survival, consistent with their role in metabolic reprogramming. In eCCA, the analysis identified a similar trend, with several EpiGs involved in chromatin regulation, such as *SETDB2* and *CHD8*, showing inverse associations with survival, in line with their upregulation in poor-prognosis groups. Regarding the MGs, no genes were consistently upregulated and negatively associated with patient survival in eCCA. Altogether, these findings highlight a set of epigenetic and metabolic regulators consistently linked to CCA aggressiveness and patient outcome across datasets.

**Table 4.**
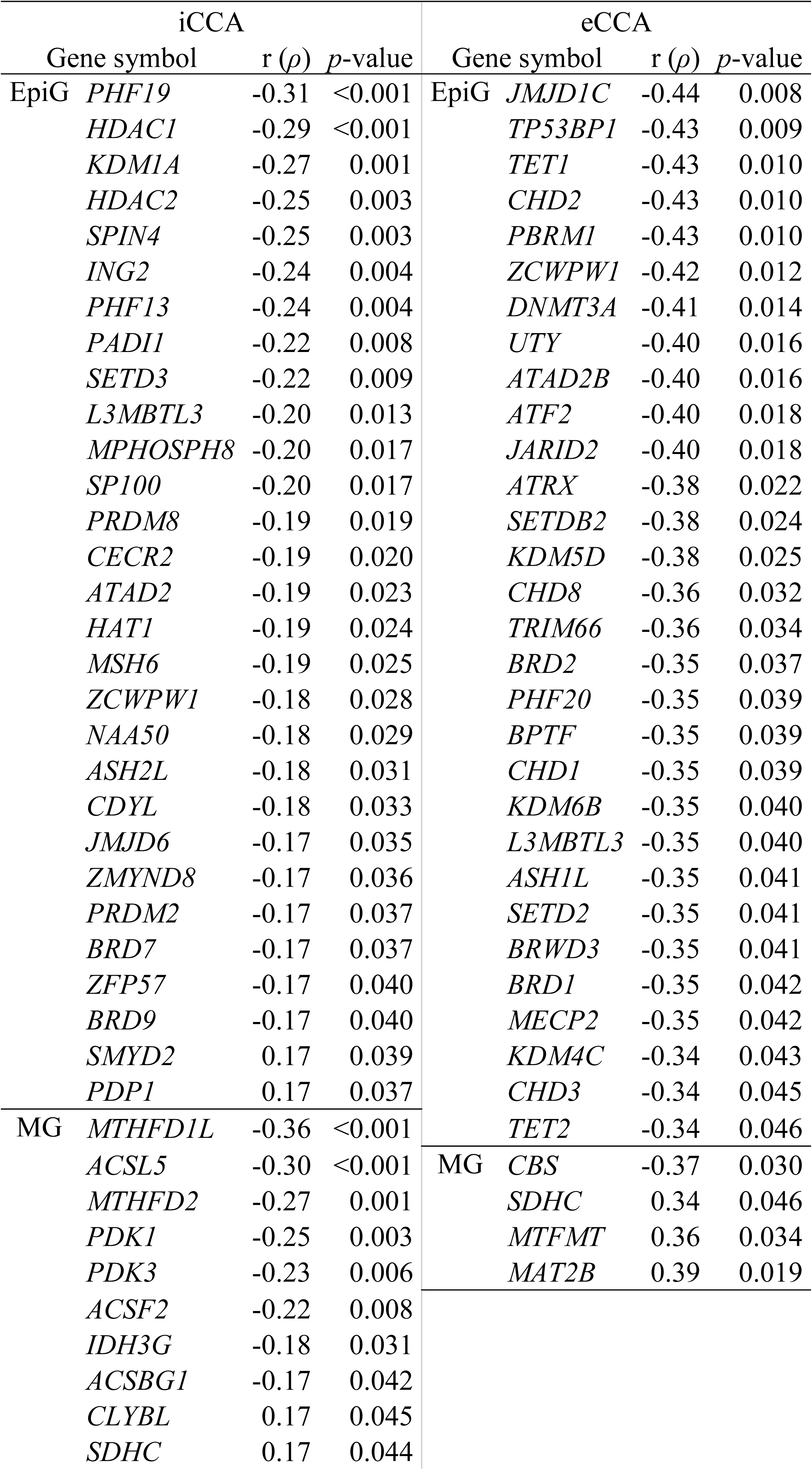
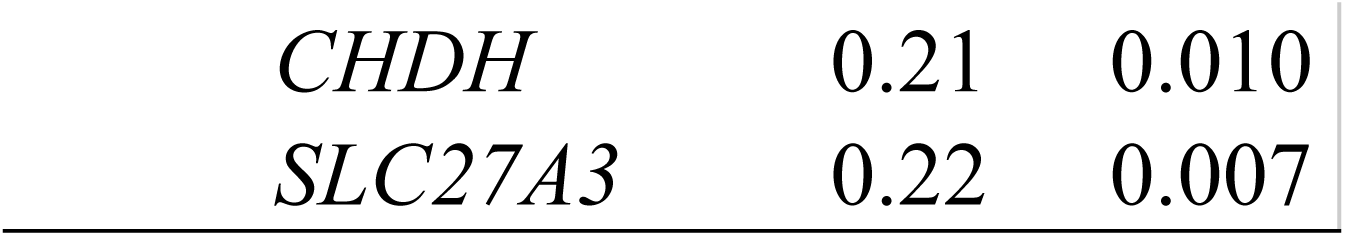
Spearman’s correlation coefficients (ρ) between the expression of epigenetic (EpiG) and metabolic (MG) genes and patient survival in the integrated CCA dataset, stratified by anatomical subtype.

| iCCA |  |  |  | eCCA |  |  |  |
| --- | --- | --- | --- | --- | --- | --- | --- |
| Gene symbol | | r ($\rho$) | p-value | Gene symbol | | r ($\rho$) | p-value |
| EpiG | *PHF19* | -0.31 | <0.001 | EpiG | *JMJD1C* | -0.44 | 0.008 |
|  | *HDAC1* | -0.29 | <0.001 |  | *TP53BP1* | -0.43 | 0.009 |
|  | *KDM1A* | -0.27 | 0.001 |  | *TET1* | -0.43 | 0.010 |
|  | *HDAC2* | -0.25 | 0.003 |  | *CHD2* | -0.43 | 0.010 |
|  | *SPIN4* | -0.25 | 0.003 |  | *PBRM1* | -0.43 | 0.010 |
|  | *ING2* | -0.24 | 0.004 |  | *ZCWPW1* | -0.42 | 0.012 |
|  | *PHF13* | -0.24 | 0.004 |  | *DNMT3A* | -0.41 | 0.014 |
|  | *PADI1* | -0.22 | 0.008 |  | *UTY* | -0.40 | 0.016 |
|  | *SETD3* | -0.22 | 0.009 |  | *ATAD2B* | -0.40 | 0.016 |
|  | *L3MBTL3* | -0.20 | 0.013 |  | *ATF2* | -0.40 | 0.018 |
|  | *MPHOSPH8* | -0.20 | 0.017 |  | *JARID2* | -0.40 | 0.018 |
|  | *SP100* | -0.20 | 0.017 |  | *ATRX* | -0.38 | 0.022 |
|  | *PRDM8* | -0.19 | 0.019 |  | *SETDB2* | -0.38 | 0.024 |
|  | *CECR2* | -0.19 | 0.020 |  | *KDM5D* | -0.38 | 0.025 |
|  | *ATAD2* | -0.19 | 0.023 |  | *CHD8* | -0.36 | 0.032 |
|  | *HAT1* | -0.19 | 0.024 |  | *TRIM66* | -0.36 | 0.034 |
|  | *MSH6* | -0.19 | 0.025 |  | *BRD2* | -0.35 | 0.037 |
|  | *ZCWPW1* | -0.18 | 0.028 |  | *PHF20* | -0.35 | 0.039 |
|  | *NAA50* | -0.18 | 0.029 |  | *BPTF* | -0.35 | 0.039 |
|  | *ASH2L* | -0.18 | 0.031 |  | *CHD1* | -0.35 | 0.039 |
|  | *CDYL* | -0.18 | 0.033 |  | *KDM6B* | -0.35 | 0.040 |
|  | *JMJD6* | -0.17 | 0.035 |  | *L3MBTL3* | -0.35 | 0.040 |
|  | *ZMYND8* | -0.17 | 0.036 |  | *ASH1L* | -0.35 | 0.041 |
|  | *PRDM2* | -0.17 | 0.037 |  | *SETD2* | -0.35 | 0.041 |
|  | *BRD7* | -0.17 | 0.037 |  | *BRWD3* | -0.35 | 0.041 |
|  | *ZFP57* | -0.17 | 0.040 |  | *BRD1* | -0.35 | 0.042 |
|  | *BRD9* | -0.17 | 0.040 |  | *MECP2* | -0.35 | 0.042 |
|  | *SMYD2* | 0.17 | 0.039 |  | *KDM4C* | -0.34 | 0.043 |
|  | *PDP1* | 0.17 | 0.037 |  | *CHD3* | -0.34 | 0.045 |
| MG | *MTHFD1L* | -0.36 | <0.001 |  | *TET2* | -0.34 | 0.046 |
|  | *ACSL5* | -0.30 | <0.001 | MG | *CBS* | -0.37 | 0.030 |
|  | *MTHFD2* | -0.27 | 0.001 |  | *SDHC* | 0.34 | 0.046 |
|  | *PDK1* | -0.25 | 0.003 |  | *MTFMT* | 0.36 | 0.034 |
|  | *PDK3* | -0.23 | 0.006 |  | *MAT2B* | 0.39 | 0.019 |
|  | *ACSF2* | -0.22 | 0.008 |  |  |  |  |
|  | *IDH3G* | -0.18 | 0.031 |  |  |  |  |
|  | *ACSBG1* | -0.17 | 0.042 |  |  |  |  |
|  | *CLYBL* | 0.17 | 0.045 |  |  |  |  |
|  | *SDHC* | 0.17 | 0.044 |  |  |  |  |

|  |  |  |
| --- | --- | --- |
| *CHDH* | 0.21 | 0.010 |
| *SLC27A3* | 0.22 | 0.007 |

### 3.5 Microenvironment-driven transcriptional programs in human CCA

We quantified immune and stromal cell populations in GSE32225 (Sia et al., 2013) using MCP-counter and derived composite scores for adaptive immunity (T cells, CD8 T cells, Cytotoxic lymphocytes, B lineage), innate immunity (NK cells, Monocytic lineage, Myeloid dendritic cells, Neutrophils), and stromal cells (Fibroblasts, Endothelial cells). These three-axis profiles were used for unsupervised clustering, which resolved four patient subtypes with distinct immune-stromal signatures: Immunogenic, Myeloid, Immune Desert, and Mesenchymal (**Figure 6A**), as previously described for CCA transcriptomics (Job et al., 2020; Ruffolo et al., 2022; Martin-Serrano et al., 2023; Zhang et al., 2023; Zhong et al., 2024). We then assessed their association with the three aggressiveness signatures reported in Sia *et al*. (Sia et al., 2013), namely proliferation-inflammation, survival, and recurrence. The Immunogenic class displayed high innate and adaptive immune scores, with moderate stromal activation, and ∼18% of iCCA patients within this class fell into poor-prognosis signature groups (**Figure 6B**). The Myeloid-rich class showed moderate-to-strong innate and adaptive immune signatures alongside low stromal scores, and comprised ∼41% patients with poor prognosis. The Immune Desert class was defined by minimal TME activity, with notably absent adaptive immunity, and comprised ∼78% of poor-prognosis patients. Lastly, the Mesenchymal class, marked by strong stromal activation and low-to-moderate immune signatures, included ∼65% poor-prognosis patients. When considering the distribution of all poor-prognosis patients across clusters, ∼5% were classified as Immunogenic, ∼21% as Myeloid-rich, ∼36% as Immune Desert, and ∼38% as Mesenchymal class.

**Figure 6.**
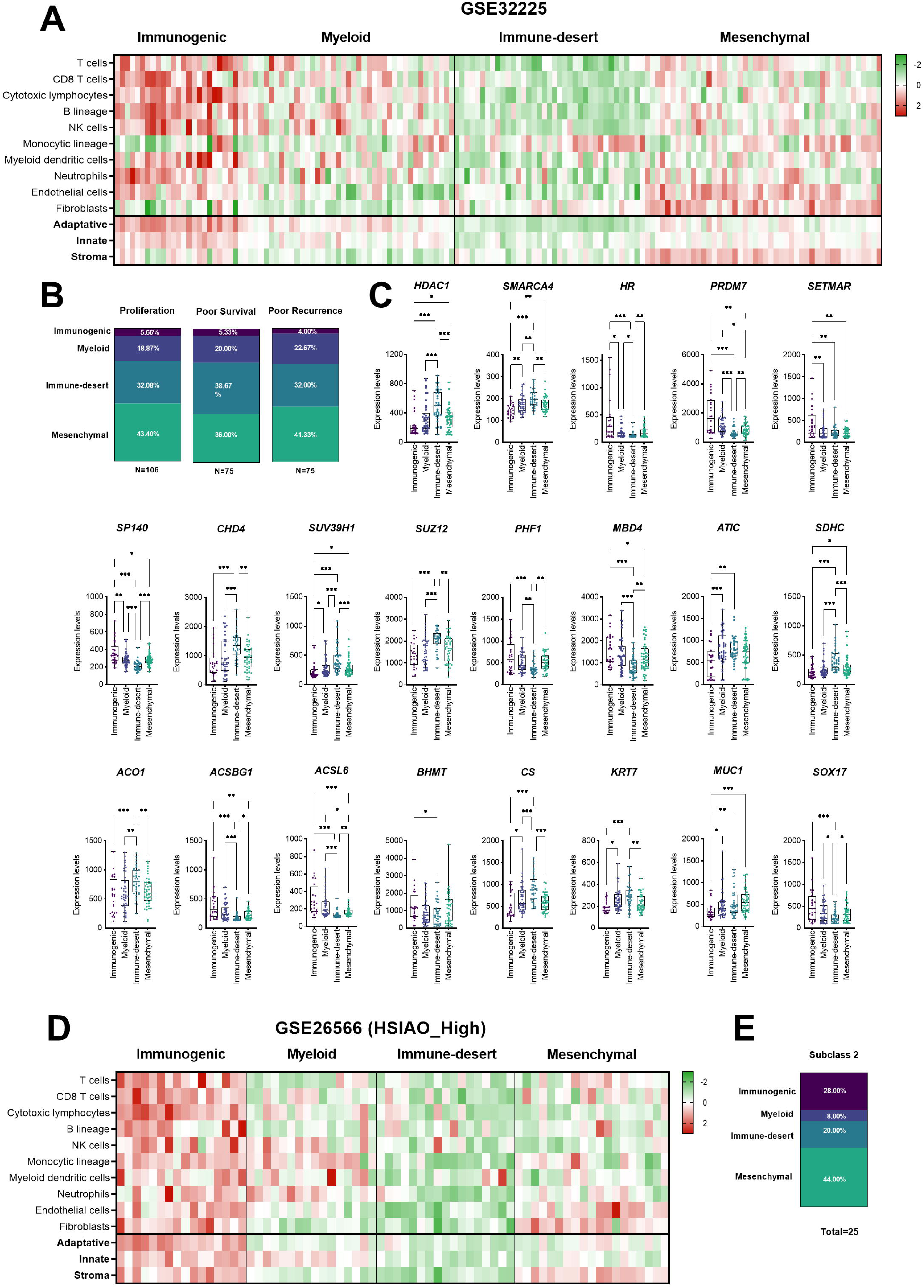
A tumor microenvironment classification of CCA patients into four immune-stromal subtypes shows deregulated epigenetic programs. **(A)** Heatmap showing MCP-counter ssGSEA enrichment scores for immune and stromal cell populations (Z-scores) in iCCA samples from GSE32225. The heatmap displays averaged Z-scores for individual immune cell types and for aggregated innate, adaptive, and stromal signatures. iCCA samples were clustered into four immune-stromal subtypes. **(B)** Proportion of iCCA patients in poor-prognosis groups from GSE32225 across the four immune-stromal clusters. **(C)** Expression levels of EpiGs, MGs, and CCA markers GSE32225 iCCA patients across immune-stromal clusters. **(D)** MCP-counter-based immune and stromal Z-score profiling of HSIAO_High (predicted iCCA) samples from GSE26566, with clustering into four immune-stromal subtypes. **(E)** Distribution of HSIAO_High (predicted iCCA) patients from GSE26566 in poor-prognosis groups across the immune-stromal clusters.

Next, by systematically selecting DEGs between clusters (raw p-value<0.05; |Log_2_FC| >0.50), we identified marked, cluster-specific transcriptional programs across EpiGs, MGs, and RLEs (complete DEGs with raw p-value<0.05 are in **Supplementary Table 10**). Several EpiGs, including *BRD2*, *CBX5*, *DEK*, *HDAC1*, *HDAC2*, *KAT2A*, *LRWD1*, *PHF20L1*, and *SMARCA4,* were upregulated, whereas *AIRE*, *BRDT*, *HR*, *PRDM7*, *SETMAR*, *SMYD1*, *SP140,* and *SPIN2A* were downregulated in the Myeloid-rich, Immune-desert, and Mesenchymal class compared with the Immunogenic class. These patterns show matching genes between the immune-stromal clusters and the CCA vs. NBD, CCA aggressiveness, and association with survival time, further supporting the Immunogenic class as the best-prognosis, immune-inflamed subtype. Notably, the Immune-desert class displayed a unique upregulation of *CHD4*, *MSH6*, *SPIN1*, *SUV39H1*, *SUZ12*, and *ZBTB33*, and downregulation of *PHF1*, consistent with a chromatin-repressive, DNA repair-oriented profile. Moreover, *HMGB1* and *SPIN4* were jointly upregulated and *MBD4* and *SETD4* downregulated in both Immune-desert and Mesenchymal classes, suggesting shared epigenetic vulnerabilities linked to stromal activation and immune exclusion. A similar pattern emerged in MGs: *AHCY*, *ATIC*, *SDHC*, and *SHMT2* were upregulated in all non-Immunogenic classes, whereas *ACSBG1*, *ACSL6*, and *IDH3A* were consistently downregulated. The Immune-desert class showed selective upregulation of *ACO1*, *MDH1*, *PDHB*, and *SDHB*, and downregulation of *BHMT*, indicating a shift toward mitochondrial and oxidative metabolic reprogramming. *CS* was elevated in both Myeloid-rich and Immune-desert classes, while *SLC27A1* was downregulated in the Immune-desert and Mesenchymal classes, suggesting reduced fatty acid uptake in immunosuppressed/stroma-dominant profiles. A selection of deregulated EpiGs and MGs across the immune-stromal clusters is shown in **Figure 6C**. RLEs also stratified classes. *ALDH2*, *HMGCS1*, *PNPO*, *PTS*, *SCD*, *SRM*, and *UGDH* were upregulated across all non-Immunogenic classes, whereas *ALDH1A2*, *CHAT*, *GCK*, *GGT1*, *LTC4S*, and *NMNAT2* were consistently downregulated. The Immune-desert class again showed the most distinctive profile, with exclusive upregulation of *ACO1*, *MDH1*, *NT5C3A,* and downregulation of *CPT1B* and *GFPT2*. Both the Myeloid-rich and Immune-desert classes exhibited downregulation of *HSD17B6* and *HDC*, and upregulation of *ODC1*, while *DDC* was reduced in Immune-desert and Mesenchymal classes. Very few genes were uniquely deregulated within a single cluster (aside from those specific to the Immune-desert group), such as the endothelial-associated RLEs *PLAT,* uniquely upregulated in the Mesenchymal class. We applied the same immune-stromal clustering strategy to the GSE26566 cohort (Andersen et al., 2012). In contrast to the iCCAs from GSE32225 (Sia et al., 2013), we observed that a substantial fraction of patients within the Immunogenic class were classified as Subclass 2, the group associated with the poorest prognosis and highest aggressiveness in this dataset (data not shown). Conversely, only a small proportion of Subclass 2 patients fell into the Immune-desert class, despite its low immune infiltration profile. This inversion of the expected “hot” (immune-inflamed) versus “cold” (immune-desert) tumor behavior, where immune-rich tumors typically show better responsiveness and more favorable outcomes, suggests that the mixed anatomical origin of CCA tumors (iCCA and eCCA) in this cohort could confound the relationship between immune microenvironment and prognosis. To minimize this source of heterogeneity, we next repeated the immune-stromal clustering exclusively in the subset of patients we reclassified as putative iCCA using the Hsiao liver-specific signature (previously mentioned as Hsiao_High group as predicted iCCAs, **Supplementary Table 6**). This refinement allowed us to reassess immune subtypes and their association with prognosis within a more anatomically homogeneous cohort. With this subset of Hsiao_High patients, we applied the same immune-stromal profiling and unsupervised clustering approach, which similarly identified four distinct patient subtypes (Immunogenic, Myeloid, Immune-desert, and Mesenchymal, **Figure 6D**), recapitulating the characteristic patterns of adaptive immunity, innate immunity, and stromal activation observed in the GSE32225 (Sia et al., 2013) cohort. Notably, some differences emerged compared to the iCCAs clustering from GSE32225. The Immunogenic class exhibited stronger stromal activation, the Mesenchymal class displayed slightly lower stromal activation than observed in the Sia immune-stromal analysis, and the Immune-desert showed lower expression across all three axes. Regardless of comparisons with the GSE32225 cohort, we evaluated the association between immune-stromal clustering of Hsiao_High patients from GSE26566 (Andersen et al., 2012) and the aggressiveness-associated subclass signature reported in that study (**Figure 6E**). In the Immunogenic class, 50% of patients fell into the poor-prognosis category, the Myeloid-rich class included 27% of poor-prognosis patients, the Immune-desert cluster had 13%, and the Mesenchymal class had the highest proportion, with 57% of patients in the poor-prognosis group (corresponding to 6%, 19%, 32% and 42% of all subclass 2 patients, respectively). To evaluate whether the immune-stromal subclasses recapitulated transcriptional trends observed in the GSE32225 cohort, we examined the gene expression of EpiGs, MGs, and RLEs that showed the same direction of deregulation in Hsiao_high patients from GSE26566 (raw p<0.05 without a magnitude threshold, **Supplementary table 10**). Among EpiG, *SUV39H1* was upregulated in the Immune-desert class, *HDAC1* in the Mesenchymal class, and *SMARCA4* in the Myeloid-rich and Immune-desert classes, all relative to the Immunogenic group. *MBD4* was downregulated in both the Immune-desert and Mesenchymal classes, and *SP140* was consistently decreased across the Myeloid-rich, Immune-desert, and Mesenchymal classes. For MGs, *ATIC* and *CS* were upregulated in the Immune-desert class, while *SDHC* was elevated in both the Immune-desert and Mesenchymal classes. Among RLEs, *DDC* and *HDC* were downregulated in the Immune-desert and Mesenchymal classes, *GFPT2* and *HSD17B6* were downregulated in the Immune-desert class, *ODC1* was upregulated in the Immune-desert class, *PTS* was upregulated in the Mesenchymal class, and *LTC4S* was downregulated in the Myeloid-rich and Immune-desert classes.

As an internal validation of this stratification, we evaluated canonical CCA markers typically deregulated in CCA in the GSE32225 immune-stromal clustering (**Figure 6C**). *MUC1* was elevated in the Myeloid-rich, Immune-desert, and Mesenchymal classes, and *KRT7* was increased in the Myeloid-rich and Immune-desert classes. Conversely, the Immune-desert class showed reduced *SOX17*, a tumor suppressor whose silencing is characteristic of aggressive CCA (Merino-Azpitarte et al., 2017). Together, these data reveal that each immune stromal subtype is defined not only by distinct TME compositions but also by coherent epigenetic, metabolic, and enzymatic transcriptional programs. These cluster-specific molecular signatures, particularly those enriched in the Immune-desert and Mesenchymal phenotypes, underscore the deep interdependence between tumor microenvironmental states and the epigenetic landscape of iCCA.

### 3.6 Epigenetic dysregulation and metabolic rewiring in experimental CCA

A preliminary indication of the potential relevance of EpiGs and MGs as therapeutic targets in CCA was obtained by assessing cell viability using gene fitness scores derived from CRISPR/Cas9 loss-of-function screens in human CCA cell lines (n = 28) from the DepMap portal. In this analysis, the individual knockout of 188 EpiGs significantly affected CCA cell fitness, with 50 genes showing absolute fitness scores above the threshold (|fitness score| >0.25) (**Figure 7A**). Several of these genes overlapped with those associated with poor prognosis or reduced patient survival, reinforcing their biological and clinical relevance. Concretely, among the EpiGs, genes such as *BRD4*, *CHD4*, *CHD8*, *DNMT1*, *HDAC3*, *MPHOSPH8*, *PRMT5*, *SETDB1*, and *SUZ12* were common to either CCA or the prognostic signatures and CRISPR-derived fitness dependencies, suggesting that these chromatin regulators are not only transcriptionally linked to aggressive CCA phenotypes but are also required for tumor cell viability. Regarding the MGs, 69 genes significantly altered cell fitness, 23 of which met the |fitness score| >0.25 criterion (**Figure 7B**). Specifically, genes such as *FH, GART, MAT2A, OGDH, SDHB, SDHC,* and *TYMS* displayed both prognostic or survival associations and significant loss-of-fitness effects upon knockout. Together, these convergent findings highlight a subset of epigenetic and metabolic regulators whose expression correlates with CCA progression and whose functional disruption compromises tumor cell survival, underscoring their potential as candidate therapeutic targets. Notably, none of the affected genes with an absolute fitness score above 0.25 showed increased cell viability; all deletions resulted in decreased fitness.

**Figure 7.**
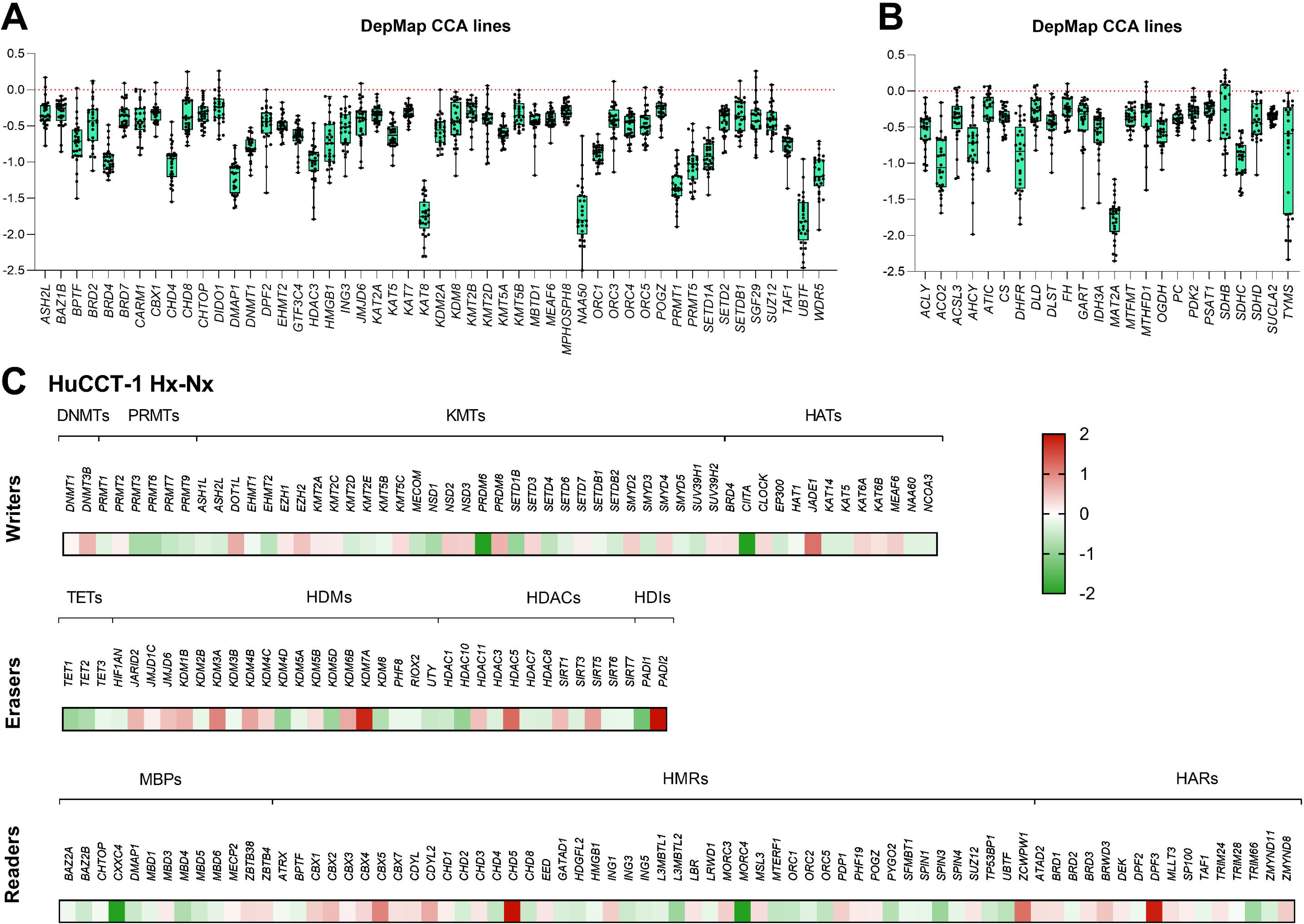
Functional dependencies and hypoxia-driven epigenetic rewiring in experimental CCA. Selected **(A)** epigenetic regulators and **(B)** metabolic genes showing gene dependency (fitness) scores in 28 CCA cell lines by CRISPR-Cas9 loss-of-function screens from DepMap. Negative scores indicate reduced viability upon gene knockout. **(C)** Expression changes of epigenetic regulators in HuCCT-1 cells exposed to 1% O₂ for 24 h, showing hypoxia-induced up-or downregulation relative to normoxia.

To further explore the mechanisms underlying epigenetic and metabolic alterations in CCA, we performed complementary experiments in human CCA cell cultures and mouse models of the disease. Since hypoxia-related pathways were consistently overrepresented across all GSEA when comparing poor-and favorable-prognosis groups in CCA, we next examined the transcriptional response of epigenetic regulators to hypoxic stress (**Figure 7C**). The human iCCA cell line HuCCT-1 was exposed to 1% O₂ for 24 h, and changes in EpiGs expression were assessed relative to normoxic conditions. Under hypoxia, 73 EpiGs were upregulated and 92 were downregulated (raw p<0.05). Notably, several genes showed concordant upregulation in hypoxic HuCCT-1 cells and in at least two of the three patient datasets comparing CCA vs. NBD, including *ATAD2*, *BRD4*, *CBX3*, *CBX4*, *DEK*, *DNMT1*, *EZH2*, *KMT2C*, *MSL3*, *NSD3*, *SP100*, and *SUZ12*, suggesting that hypoxia may contribute to the activation of a shared epigenetic program associated with malignant transformation and poor prognosis.

In parallel, we examined in vitro grown CCA tumoroids and healthy liver-derived organoids derived from patient samples, as described by Broutier et al. (Broutier et al., 2017). Because of the limited number of replicates to account for human heterogeneity, we explored EpiGs showing a raw *p*-value<0.25 and an absolute Log_2_FC>0.10 in the comparison between CCA tumoroids and healthy liver-derived organoids to identify potential expression trends (**Supplementary Table 8**). In this analysis, we found that *BRD9*, *GATAD1*, *HDAC3*, *KAT14*, *LRWD1*, *MBD4*, *ORC2*, *SETDB1*, *SGF29*, *SPIN3*, and *TAF1* were upregulated and *KMT2E* was downregulated in CCA tumoroids and at in least one of the three human CCA vs. NBD comparisons (without contradictory changes in others). Next, we identified changes in the expression (raw p-value<0.25; |Log_2_FC| >0.10) of MGs involved in key metabolic pathways that matched at least one of the three human CCA vs. NBD comparisons. Within the genes belonging to OCM pathway, *ATIC* and *SHMT2* were upregulated. Genes associated with the TCA-cycle, including *ACO2*, *CS*, and *PDHA1*, also showed increased expression. Among ACS genes, *ACSM3* was upregulated, whereas *ACSL5* was the only MGs found to be downregulated in CCA tumoroids. Finally, when analyzing genes belonging to the RLEs, we found that *ACO2*, *NT5C3A*, *PDHA1*, and *PIK3C3* were upregulated, whereas *ALDH1A2*, *ASS1*, *GK*, and *KMO* were downregulated in CCA tumoroids (raw p-value<0.25; |Log_2_FC| >0.10) and matched at least one of the three human CCA vs. NBD comparisons. These alterations further support a metabolic shift favoring mitochondrial and anabolic pathways at the expense of normal hepatobiliary metabolic functions, consistent with the transcriptional reprogramming observed in human and murine CCA models. Although the transcriptional overlap with human CCA is partial, the CCA tumoroid model reproduces several epigenetic alterations observed in human CCA, highlighting its potential as a translational system to investigate gene regulatory mechanisms *ex vivo*.

### 3.6 Multi-omic analyses identify epigenetic activation and metabolic suppression in CCA

We next established two murine CCA models by delivering oncogenes to the liver through HTVi, yielding TAZ/Akt-driven tumors and NICD1/Akt-driven tumors spanning early to advanced stages (**Figure 8A**). As a proof-of-concept, we validated our cross-species analysis of EpiGs expression by performing SMARCA4 immunostaining of human iCCA and eCCA samples and mouse CCA genetic models. This chromatin-remodeling factor (reader) has previously been shown to be upregulated at the gene expression level in various cancers, including CCA (Zhou et al., 2021), and overexpressed at the protein level in non-liver cancers (Peng et al., 2021). Also, we observed that *SMARCA4* expression was consistently upregulated across human CCA datasets. To illustrate this pattern, SMARCA4 immunostaining showed nuclear enrichment in tumor cells, with lower but detectable nuclear signal in non-tumoral hepatocytes (representative figure in **Figure 8B** and the protein expression across human and mouse CCA samples in **Supplementary Figure 3**). This example supports the robustness of our integrative approach and highlights SMARCA4 as a potential therapeutic target in human CCA.

**Figure 8.**
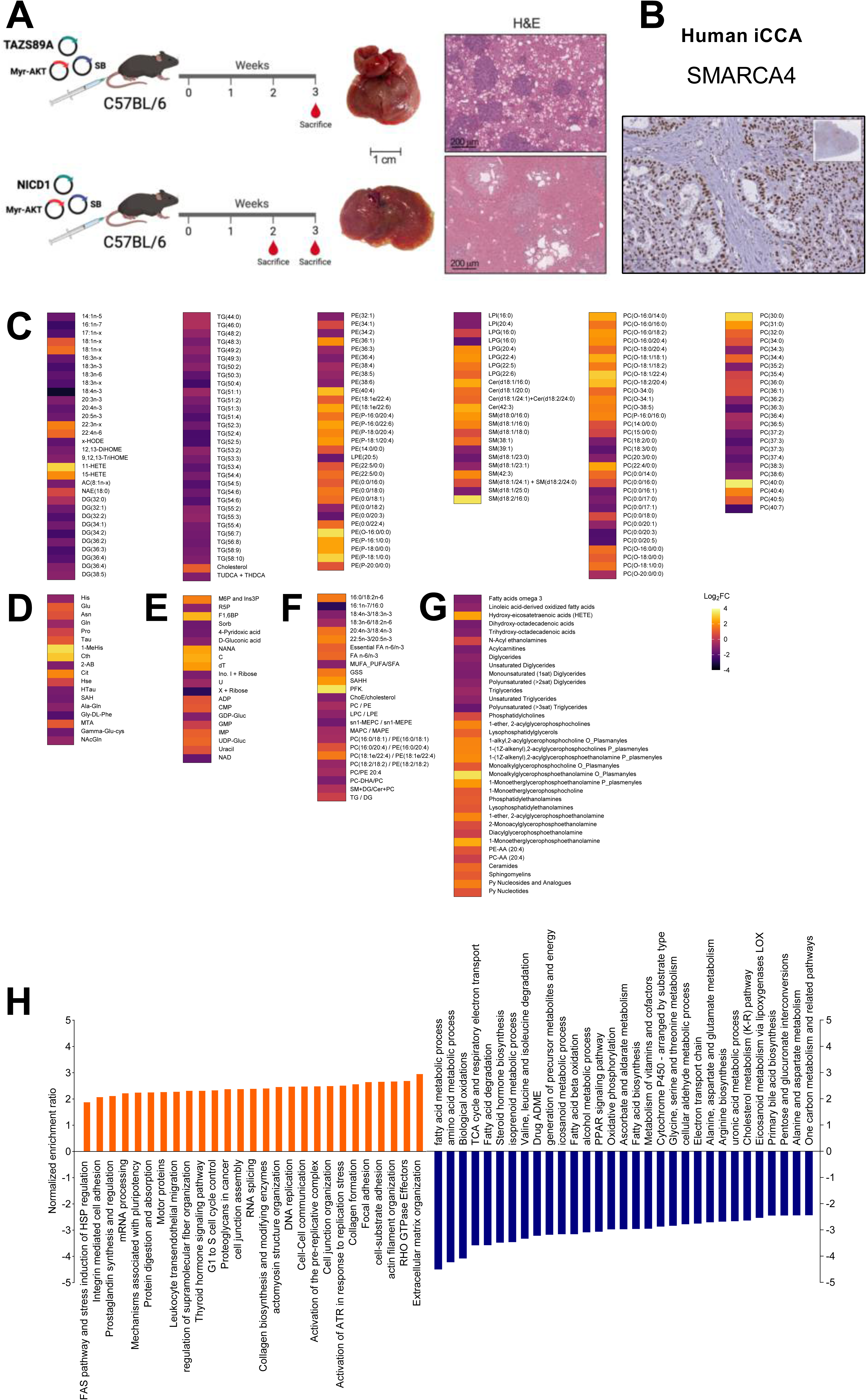
Clinical and experimental CCAs exhibit conserved epigenetic activation and metabolic reprogramming. **(A)** Generation of TAZ/Akt and NICD1/Akt murine CCA models through hydrodynamic tail-vein injection (HTVi), producing tumors spanning early to advanced disease stages (created in https://BioRender.com). **(B)** Representative image of SMARCA4 immunostaining in human iCCA, showing strong nuclear upregulation in neoplastic cells compared with adjacent non-tumoral cells. Metabolomic profiling of TAZ/Akt tumors versus healthy liver, revealing broad metabolic rewiring, including changes in **(C)** lipid classes, **(D)** amino acids, **(E)** nucleotide and carbohydrate metabolism **(F)** metabolite ratios and **(G)** metabolites grouped in families. **(H)** Pathway enrichment analysis of differentially expressed proteins in TAZ/Akt tumors.

To assess whether experimental CCA models recapitulate the transcriptional alterations observed in patients, we analyzed RNA-seq data from both human and murine systems. Transcriptomic analyses were performed in murine livers following HTVi of oncogenes, comparing normal liver tissue with tumors driven by TAZ/Akt (advanced-stage CCA) and NICD1/Akt (at early and advanced CCA stages) oncogenes. In these models, we compared the transcriptional profiles of EpiGs across human and murine datasets to identify conserved alterations in experimental CCA (**Supplementary Table 8**). We focused on EpiGs consistently deregulated in the same direction in at least two human datasets (CCA vs. NBD, raw p<0.05) and in at least one murine model (CCA vs. healthy liver, raw p<0.05): *Atad2, Cbx4, Chd4, Chd8, Dek, Dmap1, Dnmt1, Ezh2, Hdac1, Hdac2, Hdac3, Hdac7, Orc2, Prmt1, Trim28*, and *Uhrf1* were upregulated, whereas *Chd1, Ncoa1,* and *Smarca2* were downregulated. Among these, *Atad2, Chd8, Dnmt1, Ezh2, Hdac2, Hdac7, Orc2, Trim28, and Uhrf1* were increased in both genetic mouse models (TAZ/Akt and NICD1/Akt). Notably, *Atad2* and *Dnmt1* expression progressively increased during early and late tumor development in the NICD1/Akt model. In addition, several genes were upregulated in at least one human dataset (without contradictory changes in others) and in both murine models (TAZ/Akt and NICD1/Akt), including *Baz2b, Brd9, Cbx5, Ciita, Kdm2b, Kmt2a, Mbd4, Msh6, Nsd2, Orc1, Padi2, Pbrm1,* and *Pdp1*. Of particular note, *Ciita* and *Phf20,* although upregulated in only one human dataset, showed a progressive increase from early to late stages in NICD1/Akt-driven tumors. We next examined the expression of consistently deregulated MGs across human and murine models (**Supplementary Table 8**). *Aldh1l2, Idh2,* and *Tyms* were upregulated, whereas *Amt, Bhmt, Mat1a,* and *Pdk4* were downregulated in at least two human datasets (CCA vs. NBD, raw p<0.05) and in at least one murine model (CCA vs. healthy liver, raw p<0.05). Among these, *Aldh1l2* and *Idh2* were upregulated, while *Amt, Bhmt, Ftcd, Gnmt, Mthfd1,* and *Shmt1* were downregulated in both genetic mouse models (TAZ/Akt and NICD1/Akt). Particularly, *Bhmt* expression progressively decreased during early and late tumor development in the NICD1/Akt model. In addition, several MGs were deregulated in at least one human dataset (without contradictory changes in others) and in both murine models, including *Slc27a3* (upregulated) and *Acsl1, Acsl5, Acsm5, Acss3, Clybl, Glrx, Sdhd, Shmt1, Slc27a2,* and *Suclg2* (downregulated).

Given the transcriptomic evidence of coordinated epigenetic and metabolic dysregulation in human CCA, we next investigated whether these alterations were reflected at the metabolite level in mice. Metabolomic profiling was performed in the TAZ/Akt mice models recapitulating CCA features and in their healthy littermates. Thus, we evaluated differentially accumulated metabolites (DAMs), defined as metabolites that showed a statistically significant change in concentration between CCA and healthy liver tissues. DAM analysis showed broad alterations in amino acid, lipid, and nucleotide metabolism, consistent with the metabolic transcriptional reprogramming observed in human datasets (**Figure 8C-G**). Lipid metabolism displayed the most extensive remodeling (**Figure 8C**). The most strongly downregulated metabolites were the omega-3 and omega-6 PUFAs (e.g., 18:4n-3, 20:5n-3, 22:6n-3), glycerides spanning DG and TG species (e.g., DG(34:2), TG(50:4), TG(54:6)), and phospholipids such as PC and PE variants (e.g., PC(36:4), PC(38:5), PE(36:2)), together with marked decreases in oxidized linoleic acid derivatives (e.g., 9-HODE, 13-HODE, DiHOME) and selected sphingolipids (e.g., Cer(d18:1/16:0), SM(d18:1/18:0), HexCer(d18:1/22:0)). Conversely, the most upregulated metabolites included several saturated and monounsaturated PCs and PEs (e.g., PC(32:0), PC(34:1), PE(36:1)), hydroxy-eicosatetraenoic acids (HETE) species (e.g., 12-HETE, 15-HETE, 5-HETE), cholesterol, ether-and plasmalogen-linked phospholipids (e.g., PC(O-36:2), PE(P-38:4), PC(P-34:1)), and ceramides (e.g., Cer(d18:1/24:1), Cer(d18:1/22:0), Cer(d18:1/24:0)), reflecting a shift toward membrane-associated and signaling lipids. Among amino acid metabolites (**Figure 8D**), glutamic acid, asparagine, proline, taurine, L-citrulline, and cystathionine were significantly increased, whereas histidine, glutamine, S-adenosylhomocysteine, hypotaurine, and several dipeptides showed reduced levels, suggesting perturbations in one-carbon, transsulfuration, and nitrogen balance pathways. In addition, elevated methylthioadenosine (MTA) levels indicated altered methionine cycle dynamics. Beyond lipid remodeling and amino acid changes, we also detected a shift in nucleotide and carbohydrate pathways, characterized by decreased nucleosides and cofactors (e.g., xanthosine, inosine, NAD) and concomitant accumulation of nucleotide monophosphates, sugar phosphates (e.g., GMP, UDP-glucose, D-mannose 6-phosphate), and sialylated intermediates such as N-acetylneuraminic acid (**Figure 8E**). Analysis of metabolite ratios provided an empirical estimation of enzyme activities, revealing coordinated shifts in key metabolic reactions (**Figure 8F**). Ratios reflecting desaturase activity, sphingomyelin synthesis, fatty acid methylation, and cholesterol esterification were generally downregulated (e.g., SCD(n-7), sn1-MEPC/sn1-MEPE, PC-DHA/total PC, LPC/LPE), indicating reduced flux through unsaturated fatty acid, phospholipid, and sterol metabolism. Conversely, ratios associated with lipogenesis, very-long-chain fatty acid elongation, glutathione synthesis, and energy metabolism were upregulated (e.g., DGAT, PE(16:0/20:4) methylation, ELOVL5 n-3, GSS, PFK), consistent with enhanced anabolic and redox activities in CCA. These findings further support a coordinated remodeling of metabolic pathways, integrating lipid desaturation, membrane composition, and energy homeostasis. These changes were reflected at the family level (**Figure 8G**), with downregulation of PUFA-TG, PUFA-DG, omega-3 fatty acids, oxidized LA derivatives, acylcarnitines, and MUFA-DG, and upregulation of MEPE, MEPC, plasmalogen PCs and PEs, diacyl-PE, NAE, LPE, SM, and ceramides, indicating active remodeling of membrane and signaling lipids. Collectively, the metabolomic data reveal a coordinated metabolic remodeling in CCA characterized by depletion of fatty acid and acylcarnitine pools, accumulation of phospholipid and sphingolipid intermediates, and altered amino acid and methyl donor metabolism. These findings are consistent with enhanced anabolic and mitochondrial activity together with suppression of oxidative and hepatobiliary-specific processes, aligning with the transcriptional signatures of metabolic reprogramming identified in human CCA.

Proteomic profiling of the TAZ/Akt-driven iCCA model revealed extensive overlap with human CCA transcriptomic data (CCA vs. NBD), supporting its translational relevance (**Supplementary Table 8**). Several epigenetic regulators (CBX3, CHD4, DEK, DNMT1, HDAC1, HDAC2, PRMT1, and TRIM28), metabolic enzymes (TYMS and IDH2), and RLEs (PKM, TKT, and TYMS) showed concordant upregulation at the mRNA and protein level across at least two human datasets and in the TAZ/Akt mouse model. In contrast, key metabolic enzymes (ALDH1L1, FTCD, GNMT, MTHFD1, and SHMT1) and RLEs genes (ACADL, ALDH2, CSP1, DDC, FBP1, HSD17B6, and LIPC) were consistently downregulated. Also, the writer BRD4 and the OCM enzyme ATIC emerged as recurrently upregulated DEP shared with human datasets at the mRNA level (CCA vs. NBD), although not transcriptionally induced in the TAZ/Akt model. Pathway enrichment analysis of DEPs (**Figure 8H)** showed a marked imbalance between metabolic functions and pathways associated with chromatin regulation and oncogenic signaling. Underrepresented terms were dominated by metabolic hubs, including central carbon and energy pathways such as citrate cycle (TCA cycle) and oxidative phosphorylation, together with amino acid and lipid catabolic programs (valine, leucine and isoleucine degradation, fatty acid metabolic process, arachidonic acid metabolism). Pathways linked to xenobiotic processing and detoxification, including drug metabolism and cytochrome P450-mediated xenobiotic metabolism, were also consistently reduced, suggesting a broad suppression of mitochondrial and peroxisomal functions. In contrast, overrepresented terms highlighted a strong activation of epigenetic and chromatin-remodeling modules, including the polycomb repressive complex, histone deacetylase complex, and ATP-dependent chromatin remodeling. These changes were accompanied by the enrichment of oncogenic signaling networks central to CCA development and malignancy, including the TGF-beta, Notch, Ras, PI3K-Akt, and Hippo signaling pathways. Together, these findings indicate that CCA progression is characterized by coordinated downregulation of metabolic competence and parallel reinforcement of chromatin-based regulatory mechanisms and proliferative signaling cascades. Together, these proteomic alterations indicate a shift from hepatocyte-specific, oxidative metabolic programs toward proliferative, adhesive, and stromal-interacting states characteristic of CCA.

## 4 Discussion

Our integrative analysis reveals an extensive rewiring of both epigenetic regulators and metabolism in CCA, highlighting the interdependence of epigenetic regulation and cellular metabolic states in driving tumor progression. Across CCA patient cohorts, we observed consistent upregulation of epigenetic writers, erasers, and readers, including *HDAC3*, *PHF20L1*, *CBX3*, *EZH2,* and *SMARCA4*, alongside downregulation of readers such as *CHD5* in all CCA datasets when compared to healthy tissues (NBD). Several of these upregulated EpiGs, such as HDAC3 and EZH2, have been previously studied in the context of CCA (Yin et al., 2017; Wasenang et al., 2019; Zhang et al., 2022; Wu et al., 2023), whereas CBX3 has been recently reported to promote proliferation and invasion in HuCCT-1 and RBE CCA cells (Xie et al., 2025). The readers SMARCA4, already described to promote hepatocellular carcinoma (HCC) proliferation (Chen et al., 2018) and pinpointed in pan-cancer studies (Peng et al., 2021), *PHF20L1*, found as carcinogenic in breast, ovarian, and colorectal cancer (Chen et al., 2018; Hou et al., 2020; Alberto-Aguilar et al., 2022), and CHD5, a tumor suppressor in HCC (Zhao et al., 2014), have not been described before as potential tumor-related genes in human CCA. These alterations converge on pathways related to nucleosome organization, chromatin remodeling, DNA repair, and transcriptional elongation, underscoring the establishment of an epigenetic landscape that favors proliferation, mitotic fidelity, and oncogenic transcriptional programs in CCA (O’Rourke et al., 2018, 2019; Banales et al., 2020; Zhong et al., 2023). Metabolic enzymes that generate key cofactors for epigenetic reactions (Ghisletti and Russo, 2025) were also consistently dysregulated at the transcriptomic level, with selective upregulation of OCM-and TCA-related components and suppression of enzymes involved in methyl-donor homeostasis, TCA flux, and fatty acid activation as previously described (Raggi et al., 2022; Lori et al., 2024; Natarajan et al., 2025). Homocysteine can be remethylated back into methionine via BHMT or diverted into the transsulfuration pathway via CBS, linking one-carbon metabolism with redox homeostasis (Ducker and Rabinowitz, 2017). In parallel, decarboxylation of SAM by AMD1 initiates polyamine synthesis and produces MTA as a by-product. MTA is metabolized by MTAP, which prevents its accumulation; because MTA can inhibit methyltransferases, MTAP loss reshapes SAM/SAH/MTA homeostasis and contributes to epigenetic dysregulation and cancer progression (Feron, 2019). Complementing these methylation-dependent pathways, the availability of acetyl-CoA and other acyl-CoAs critically determines histone acetylation and lysine acylation. In cancer cells, acetyl-CoA is supplied by PDH, ACSS2, and ACLY, while TCA cycle enzymes such as OGDH and the BCKDH complex generate additional acyl-CoA species that support non-acetyl acylations (Jo et al., 2020; Guertin and Wellen, 2023; Ghisletti and Russo, 2025). Dysregulation of these interconnected routes reshapes both methyl-donor and acyl-CoA pools, altering chromatin-modifying reactions and contributing to the epigenetic plasticity (Jo et al., 2020) characteristic of CCA. In parallel, α-KG-dependent demethylases are influenced by TCA-cycle enzymes such as FH, SDHA/B, ACO2, and IDH1/2, which modulate α-KG and oncometabolite levels (Yang et al., 2013; Wang and Lei, 2018). Disruptions in these interconnected pathways collectively reshape methyl-donor, acyl-CoA, and α-KG availability (Ghisletti and Russo, 2025), driving the epigenetic plasticity characteristic of CCA. Broader pathway analysis confirmed a metabolic shift characterized by the loss of amino acid, lipid, and xenobiotic metabolism and by the enrichment of oxidative phosphorylation, glycolysis, and proteostatic programs, consistent with increased biosynthetic demand. Chromatin modifiers depend on metabolites (SAM, acetyl-CoA, NAD⁺, α-KG) whose levels are set by pathway flux (Boon, 2021; Ghisletti and Russo, 2025); thus, alterations in RLEs expose metabolic bottlenecks that impact cofactor availability and influence epigenetic regulation in CCA (Boon et al., 2020). In our analyses, deregulated RLEs and pathways further confirmed the induction of anabolic and redox-supporting factors and suppression of fatty acid oxidation, gluconeogenesis, and hepatobiliary functions (Raggi et al., 2022). Together, these transcriptomic changes delineate a dedifferentiated metabolic state that supports and reinforces the epigenetic programs maintaining malignant cholangiocyte identity.

Importantly, epigenetic dysregulation was associated with clinical aggressiveness, reflected in the consistent upregulation of *CHD4*, *CHD8*, *CIITA*, *HDAC1*, *HDAC2*, *MECOM*, *MPHOSPH8*, *PRDM8*, *PRMT1*, *SETDB2*, *SP100*, *SPIN4*, and *TDRD3* in aggressive subclasses, and the parallel downregulation of *BPTF*, *DNMT3L*, *ING5*, *LBR*, *MEAF6*, *MORC4*, *SETMAR*, and *SPIN2A*. The GSEA indicates that the dysregulation of EpiGs in aggressive CCA subclasses converges on programs controlling proliferative signaling, mitotic progression, and transcriptional remodeling, consistent with the establishment of an epigenetic landscape that favors tumor growth and poor clinical outcome. Consistent with epigenetic alterations, MGs in aggressive CCA subclasses showed recurrent upregulation of OCM and lipid metabolism enzymes (e.g., *AHCYL2*, *ACSL5, SHMT2*) and downregulation of mitochondrial and oxidative components (e.g., *ACSL6*, *CHDH, OGDHL*). These patterns were confirmed across prognostic groups, proliferation/inflammation subclasses, and recurrence analyses, and discriminant MGs were identified by DAPC. We have previously shown that several of these EpiGs were also dysregulated in other liver disease and liver cancer contexts (Clavería-Cabello et al., 2023; Herranz et al., 2023; Castelló-Uribe et al., 2025). Together, the results indicate coordinated metabolic reprogramming that complements epigenetic dysregulation, also associated with poor clinical outcomes. In fact, several genes showed inverse association with the iCCA patient’s survival in a validation cohort (e.g., *HDAC1*, *HDAC2*, *SPIN4*, *PRDM8*, *MSH6*). Some of these epigenetic regulators, such as *HDAC1* (Xu et al., 2022; Xiong et al., 2024) and *HDAC2* (He et al., 2016), have already been associated with worse patient prognosis and tumor aggressiveness in CCA. However, some epigenetic regulators have a hitherto unknown role in CCA.

The MCP-counter method allowed us to quantify the abundance of immune and stromal cell populations in human CCA tissue samples using transcriptomic data, providing insight into the epigenetic and metabolic landscape within the TME. The cell population scores were then grouped into adaptive, innate, and stromal scores to represent the tumor microenvironment. Although *in silico* gene expression deconvolution has already been employed for patient stratification in CCA (Job et al., 2020; Montal et al., 2020; Martin-Serrano et al., 2023), the epigenetic landscape of the resulting immune-stromal subtypes has not yet been delineated. An unsupervised clustering was applied to these adaptive, innate, and stromal scores, further stratifying CCA into distinct immune-stromal subtypes (Immunogenic, Myeloid-rich, Immune-desert, and Mesenchymal classes), each characterized by coherent epigenetic and metabolic programs. The Immunogenic class, marked by robust adaptive and innate immune activity, aligns with lower aggressiveness. In contrast, Immune-desert and Mesenchymal classes exhibit chromatin-repressive signatures, mitochondrial and oxidative metabolic shifts, and stromal activation, all of which are associated with poorer prognosis. Notably, lactate accumulation in the TME, resulting from tumor glycolysis, has been shown to modulate immune cell activity and epigenetic programs, contributing to immune exclusion and metabolic rewiring in aggressive CCA subtypes(Dong et al., 2025). Several epigenetic and metabolic regulators showed consistent deregulation across the immune-stromal patient clusters and independent datasets, reinforcing their potential role in shaping iCCA biology. *SUV39H1* was uniquely upregulated in the Immune-desert subtype, in line with its increased expression in GSE32225 (iCCA vs. NBD) and in the TAZ/Akt mouse CCA model, suggesting a link between repressive chromatin states and immune exclusion. *HDAC1* was also elevated in the Immune-desert class, consistent with its known overexpression in human CCA and further validated across GSE32225, GSE26566, and the TAZ/Akt model at both RNA and protein levels. *SMARCA4* was upregulated in the Myeloid-rich, Immune-desert, and Mesenchymal subtypes, mirroring its increased expression in CCA relative to NBD in GSE32225, GSE26566, and GSE132305, and in Hsiao_High-reclassified samples, supporting a broad role in CCA chromatin remodeling. *MBD4*, increased in Immune-desert and Mesenchymal subtypes, was also elevated in GSE132305 (eCCA vs. NBD), human tumoroids, and both TAZ/Akt and NICD1/Akt mouse models. In contrast, *SP140* was consistently decreased in the non-Immunogenic clusters and in GSE26566, suggesting a potential association with the loss of immune-inflamed phenotypes. Among metabolic genes, *ATIC* was strongly upregulated across multiple datasets (GSE32225, GSE26566, human tumoroids), and at the protein level in TAZ/Akt tumors. *SDHC*, elevated in all non-Immunogenic clusters, was also upregulated in GSE32225 and GSE132305, though downregulated in both mouse CCA models, pointing to potential species-specific or stage-dependent regulation. Together, these patterns highlight a convergence between immune-stromal architecture and epigenetic-metabolic reprogramming in CCA.

In the mesenchymal cluster the RLEs *PLAT* was upregulated compared to the other clusters, as expected since the encoded enzyme is involved in the fibrinolytic cascade and mainly expressed in endothelial cells, recently suggested as a proliferation and invasion mediator in CCA due to its relation to galectin-1 (Sirica et al., 2009). *PRMT5*, and *EHMT2*, already described to be involved in CCA and HCC (Bárcena-Varela et al., 2019; Colyn et al., 2021, 2022; Elurbide et al., 2025), although did not pass the threshold of 0.5 Log2FC, were upregulated in all groups compared to the Immunogenic group in both cohorts analyzed. These epigenetic factors have also been linked to immune regulation in other cancers, including CCA, pancreatic ductal adenocarcinoma and glioma (Cao et al., 2023; Elurbide et al., 2025; Oyon et al., 2025). These findings indicate that microenvironmental context shapes, and is shaped by, intrinsic epigenetic and metabolic states, highlighting potential vulnerabilities for targeted therapy.

Experimental models recapitulated these patient-derived patterns. Hypoxia is a well-recognized driver of tumor progression affecting practically all hallmarks of cancer (Sin et al., 2023; Liang et al., 2024; Acuña-Pilarte and Koh, 2025). However, the epigenetic dimension of hypoxia-driven tumor progression is still emerging. Interestingly, hypoxia-mediated regulation of histone demethylases such as JMJD1A (KDM3A) and KDM5A (Krieg et al., 2010; Li et al., 2021), accumulation of methylated histones and the coordinated remodeling of histone marks under low oxygen have been described (Lee et al., 2017; Chang et al., 2025). Importantly, the modulation of α-KG-dependent demethylases such as KDM5C and KDM6A by oncometabolites and oxygen availability was previously reported (Chang et al., 2019). We observed that exposure of the HuCCT-1 iCCA cell line to hypoxia induced the expression of key epigenetic regulators similarly altered in tumors. At the same time, human CCA tumoroids and murine NICD1/Akt or TAZ/Akt models partially mirrored both transcriptional and metabolic alterations observed in patients. Functional CRISPR screens reinforced this notion, demonstrating that loss of EpiGs or MGs genes associated with poor prognosis significantly reduces CCA cell fitness, validating these pathways as potential therapeutic targets. Interestingly, the magnitude of change was greater for MGs than for EpiGs when comparing CCA tumoroids to bile duct organoids. This suggests that the transition from tissue extraction to the establishment of 3D cultures exerts a stronger effect on metabolic gene expression than on epigenetic regulator expression. These findings imply that the culture and adaptation processes may preferentially alter metabolic programs, while epigenetic landscapes remain relatively more stable during *ex vivo* expansion.

Multi-omics and cross-species analyses identified conserved upregulation of methyltransferases and acetyl transferases (DNMT1, PRMT1, BRD4), histone deacetylases (HDAC1, HDAC2), and histone methyl and acetyl readers (CBX3, CHD4; TRIM28, DEK, SMARCA4) and suppression of metabolic enzymes (ATIC, IDH2, TYMS), and RLEs (PKM, TKT, TYMS), confirming that these programs are functionally relevant and evolutionarily conserved. Metabolomic profiling in TAZ/Akt-driven murine tumors mirrored these transcriptional changes, demonstrating depletion of fatty acid and acylcarnitine pools, alongside accumulation of phospholipid intermediates and alterations in amino acid and methyl donor metabolism. Pathway analyses of DEP further confirmed the coordination between metabolic suppression and epigenetic activation at the protein level, emphasizing that CCA progression entails a tightly orchestrated reprogramming of both gene expression and metabolite networks.

The recurrent dysregulation of EpiGs in both iCCA and eCCA, as well as in poor-prognosis and high-aggressiveness subclasses, indicates a central and conserved role of epigenetic modulation in CCA pathobiology. Our data also reveal clear distinctions between iCCA and eCCA, with subsets of epigenetic and metabolic regulators showing inverse expression patterns and divergent pathway enrichment profiles. iCCAs were enriched for stress response and signaling pathways, while eCCAs retained lipid, detoxification, and extracellular matrix-related programs. The divergent expression patterns of EpiGs between iCCA and eCCA suggest that each subtype engages distinct epigenetic and metabolic programs, shaped by their specific cellular origins, etiologic contexts, and microenvironmental cues that differentiate intrahepatic from extrahepatic biliary epithelia. At the same time, discrepancies between datasets likely reflect the distinct nature of the control samples beyond the intrinsic biological differences between iCCA and eCCA. For instance, in the Montal et al. study (Montal et al., 2020), the NBD corresponds to peritumoral bile duct tissue, which may influence gene expression patterns. Similarly, in other cancers such as gliomas, recent studies have highlighted how tumor-intrinsic and microenvironmental metabolic cues, including SAM and α-KG, regulate chromatin states, illustrating that the dynamic interplay between metabolism and epigenetics is a broader mechanism influencing tumor biology and therapeutic opportunities (Natarajan et al., 2025).

Collectively, our findings highlight that CCA is characterized by a coordinated rewiring of epigenetic and metabolic networks that sustains proliferation, supports survival under microenvironmental stress, and drives aggressive tumor behavior. This integrated epigenetic-metabolic landscape provides mechanistic insight into tumor heterogeneity, identifies conserved vulnerabilities, and offers a framework for future therapeutic interventions targeting both chromatin regulation and metabolic adaptation. In fact, there are already approved epidrugs that modulate the activity of a growing number of epigenetic effectors such as *HDAC3, BRD4, CIITA, DNMT1, EHMT2, EZH2, HDAC1, HDAC2, KDM5C, PRMT1, PRMT5, SETDB2, SMARCA4,* and *SUV39H1*, and some of them are undergoing clinical trials either alone or in combination with chemo-and immunetherapies for solid tumors (Bueloni et al., 2025; Fernandez-Barrena et al., 2025; Khan et al., 2025; Talom et al., 2025).

## 5 Conclusions

Our integrative analyses show that CCA is characterized by a coordinated reorganization of epigenetic regulators and metabolic pathways. Alterations in EpiGs and MGs are consistent across patient cohorts, experimental models, and aggressive molecular subclasses, linking chromatin plasticity to metabolic adaptation that supports proliferation, stress resilience, and tumor progression. While iCCA and eCCA share a common epigenetic-metabolic framework, subtype-specific differences reflect the influence of cellular origin, microenvironment, and metabolic context. Transcriptomic, proteomic and metabolomic profiling in the TAZ/Akt mouse CCA model confirmed the conservation of these programs observed in human CCAs at a multi-omic level. In parallel, deconvolution analysis of the TME in human CCA datasets allowed us to *in silico* evaluate the epigenetic landscape across immune-stromal subtypes, revealing how chromatin states intersect with microenvironmental composition. Functional validation identifies previously unrecognized CCA drivers, highlighting their potential as therapeutic targets. These findings provide mechanistic insight into tumor heterogeneity, expose vulnerabilities for epigenetic-and metabolism-directed interventions, and offer a framework for prioritizing candidate targets in precision therapies for CCA.

## Author Contributions

Conceptualization and study design were performed by A.L-P., M.A.A. and M.G.F-B.; human CCA data search, integration and analyses: A.L-P., B.C-U., E.A., M.A.A. and M.G.F-B.; Experimental models and analyses: J.E., E.V-G., M.U.L., E.A-V., L.A.M-P., I.U., S.C., F.J.C., S.F., E.S., O.F., L.C., P.I., R.A-B., Manuscript draft: A.L-P., M.A.A. critical review and editing: A.L-P., M.A., J.B., M.H., C.B., M.G.F-B., M.A.A. All authors have read and approved the final manuscript.

## 9 Acknowledgments

The technical support of Mr Roberto Barbero and Ms Miriam Belzunce is acknowledged. The authors are grateful to Mr Eduardo Avila for his geneorus sponsorship.

## 11 Supplementary Material

The Supplementary Material for this article can be found online.

## 12 Data Availability Statement

All human CCA microarray datasets used in this study, including GSE32225, GSE26566, and GSE132305, are publicly available through the NCBI Gene Expression Omnibus (GEO). Human CCA RNA-seq datasets can be accessed from their respective repositories: TCGA-CHOL (phs000178) gene expression data available at the TCGA research network (https://www.cancer.gov/tcga), and EGAD00001001693 available via the European Genome-phenome Archive (EGA, https://ega-archive.org/). RNA-seq data generated from the HuCCT-1 CCA cell line under hypoxic conditions (1% O₂) have been deposited in the European Nucleotide Archive (ENA) under accession number RJEB101122. High-throughput data on the TAZ/Akt model are available on reasonable request.

## References

Acuña-Pilarte, K., and Koh, M. Y. (2025). The HIF axes in cancer: angiogenesis, metabolism, and immune-modulation. Trends Biochem. Sci. 50, 677–694. doi:10.1016/j.tibs.2025.06.005.

Ahn, K. S., O’Brien, D., Kang, Y. N., Mounajjed, T., Kim, Y. H., Kim, T. S., et al. (2019). Prognostic subclass of intrahepatic cholangiocarcinoma by integrative molecular-clinical analysis and potential targeted approach. Hepatol. Int. 13, 490–500. doi:10.1007/S12072-019-09954-3.

Akce, M., El-Rayes, B. F., and Wajapeyee, N. (2023). Combinatorial targeting of immune checkpoints and epigenetic regulators for hepatocellular carcinoma therapy. 42, 1051–1057. doi:10.1038/S41388-023-02646-1.

Alberto-Aguilar, D. R., Hernández-Ramírez, V. I., Osorio-Trujillo, J. C., Gallardo-Rincón, D., Toledo-Leyva, A., and Talamás-Rohana, P. (2022). PHD finger protein 20-like protein 1 (PHF20L1) in ovarian cancer: from its overexpression in tissue to its upregulation by the ascites microenvironment. Cancer Cell Int. 22, 6. doi:10.1186/s12935-021-02425-6.

Ally, A., Balasundaram, M., Carlsen, R., Chuah, E., Clarke, A., Dhalla, N., et al. (2017). Comprehensive and Integrative Genomic Characterization of Hepatocellular Carcinoma. Cell 169, 1327–1341.e23. doi:10.1016/j.cell.2017.05.046.

Alvaro, D., Gores, G. J., Walicki, J., Hassan, C., Sapisochin, G., Komuta, M., et al. (2023). EASL-ILCA Clinical Practice Guidelines on the management of intrahepatic cholangiocarcinoma. J. Hepatol. 79, 181–208. doi:10.1016/J.JHEP.2023.03.010.

Anders, S., Pyl, P. T., and Huber, W. (2015). HTSeq-A Python framework to work with high-throughput sequencing data. Bioinformatics 31, 166–169. doi:10.1093/bioinformatics/btu638.

Andersen, J. B., Spee, B., Blechacz, B. R., Avital, I., Komuta, M., Barbour, A., et al. (2012). Genomic and Genetic Characterization of Cholangiocarcinoma Identifies Therapeutic Targets for Tyrosine Kinase Inhibitors. Gastroenterology 142, 1021. doi:10.1053/J.GASTRO.2011.12.005.

Banales, J. M., Marin, J. J. G., Lamarca, A., Rodrigues, P. M., Khan, S. A., Roberts, L. R., et al. (2020). Cholangiocarcinoma 2020: the next horizon in mechanisms and management. Nat. Rev. Gastroenterol. Hepatol. 17, 557–588. doi:10.1038/s41575-020-0310-z.

Bao, X., Li, Q., Chen, J., Chen, D., Ye, C., Dai, X., et al. (2022). Molecular Subgroups of Intrahepatic Cholangiocarcinoma Discovered by Single-Cell RNA Sequencing-Assisted Multiomics Analysis. Cancer Immunol. Res. 10, 811–828. doi:10.1158/2326-6066.CIR-21-1101.

Bárcena-Varela, M., Caruso, S., Llerena, S., Álvarez-Sola, G., Uriarte, I., Latasa, M. U., et al. (2019). Dual Targeting of Histone Methyltransferase G9a and DNA-Methyltransferase 1 for the Treatment of Experimental Hepatocellular Carcinoma. Hepatology 69, 587–603. doi:10.1002/hep.30168.

Barr, J., Caballería, J., Martínez-Arranz, I., Domínguez-Díez, A., Alonso, C., Muntané, J., et al. (2012). Obesity-dependent metabolic signatures associated with nonalcoholic fatty liver disease progression. J. Proteome Res. 11, 2521–2532. doi:10.1021/PR201223P.

Bates, S. E. (2020). Epigenetic Therapies for Cancer. N. Engl. J. Med. 383, 650–663. doi:10.1056/NEJMra1805035.

Becht, E., Giraldo, N. A., Lacroix, L., Buttard, B., Elarouci, N., Petitprez, F., et al. (2016). Estimating the population abundance of tissue-infiltrating immune and stromal cell populations using gene expression. Genome Biol. 2016 *171* 17, 218-. doi:10.1186/S13059-016-1070-5.

Behan, F. M., Iorio, F., Picco, G., Gonçalves, E., Beaver, C. M., Migliardi, G., et al. (2019). Prioritization of cancer therapeutic targets using CRISPR-Cas9 screens. Nature 568, 511–516. doi:10.1038/S41586-019-1103-9.

Biswas, S., and Rao, C. M. (2018). Epigenetic tools (The Writers, The Readers and The Erasers) and their implications in cancer therapy. Eur. J. Pharmacol. 837, 8–24. doi:10.1016/j.ejphar.2018.08.021.

Boon, R. (2021). Metabolic Fuel for Epigenetic: Nuclear Production Meets Local Consumption. Front. Genet. 12, 768996. doi:10.3389/fgene.2021.768996.

Boon, R., Silveira, G. G., and Mostoslavsky, R. (2020). Nuclear metabolism and the regulation of the epigenome. Nat. Metab. 2, 1190–1203. doi:10.1038/S42255-020-00285-4.

Brindley, P. J., Bachini, M., Ilyas, S. I., Khan, S. A., Loukas, A., Sirica, A. E., et al. (2021). Cholangiocarcinoma. Nat. Rev. Dis. Prim. 7. doi:10.1038/s41572-021-00300-2.

Broutier, L., Mastrogiovanni, G., Verstegen, M. M. A., Francies, H. E., Gavarró, L. M., Bradshaw, C. R., et al. (2017). Human primary liver cancer–derived organoid cultures for disease modeling and drug screening. Nat. Med. 23, 1424–1435. doi:10.1038/nm.4438.

Bueloni, B., G Fernandez-Barrena, M., Fiore, E., Avila, M. A., Bayo, J., and Mazzolini, G. D. (2025). Epigenetic therapies in hepatocellular carcinoma: emerging clinical tools and applications. Gut. doi:10.1136/gutjnl-2025-336317.

Cao, Y., Liu, B., Cai, L., Li, Y., Huang, Y., Zhou, Y., et al. (2023). G9a promotes immune suppression by targeting the Fbxw7/Notch pathway in glioma stem cells. CNS Neurosci. Ther. 29, 2508–2521. doi:10.1111/cns.14191.

Castelló-Uribe, B., Amaya López-Pascual, Elurbide, Jasmin, Adán-Villaescusa, E., Emiliana Valbuena-Goiricelaya, Martinez-Perez, L. A., et al. (2025). Expression landscape of epigenetic genes in human hepatocellular carcinoma. J. Physiol. Biochem. 2025, 1–29. doi:10.1007/S13105-025-01095-6.

Chaisaingmongkol, J., Budhu, A., Dang, H., Rabibhadana, S., Pupacdi, B., Kwon, S. M., et al. (2017). Common Molecular Subtypes Among Asian Hepatocellular Carcinoma and Cholangiocarcinoma. Cancer Cell 32, 57–70.e3. doi:10.1016/J.CCELL.2017.05.009.

Chang, S., Moon, R., Nam, D., Lee, S.-W., Yoon, I., Lee, D.-S., et al. (2025). Hypoxia increases methylated histones to prevent histone clipping and heterochromatin redistribution during Raf-induced senescence. Nucleic Acids Res. 53. doi:10.1093/nar/gkae1210.

Chang, S., Yim, S., and Park, H. (2019). The cancer driver genes IDH1/2, JARID1C/ KDM5C, and UTX/ KDM6A: crosstalk between histone demethylation and hypoxic reprogramming in cancer metabolism. Exp. Mol. Med. 51, 1–17. doi:10.1038/s12276-019-0230-6.

Chen, J., Wang, D., Wu, G., Xiong, F., Liu, W., Wang, Q., et al. (2024). STUB1-mediated K63-linked ubiquitination of UHRF1 promotes the progression of cholangiocarcinoma by maintaining DNA hypermethylation of PLA2G2A. J. Exp. Clin. Cancer Res. 43, 260. doi:10.1186/s13046-024-03186-6.

Chen, X., Dong, L., Chen, L., Wang, Y., Du, J., Ma, L., et al. (2023). Epigenome-wide development and validation of a prognostic methylation score in intrahepatic cholangiocarcinoma based on machine learning strategies. Hepatobiliary Surg. Nutr. 12, 478–494. doi:10.21037/HBSN-21-424.

Chen, Z., Lu, X., Jia, D., Jing, Y., Chen, D., Wang, Q., et al. (2018). Hepatic SMARCA4 predicts HCC recurrence and promotes tumour cell proliferation by regulating SMAD6 expression. Cell Death Dis. 9, 59. doi:10.1038/s41419-017-0090-8.

Cigliano, A., Zhang, S., Ribback, S., Steinmann, S., Sini, M., Ament, C. E., et al. (2022). The Hippo pathway effector TAZ induces intrahepatic cholangiocarcinoma in mice and is ubiquitously activated in the human disease. J. Exp. Clin. Cancer Res. 41, 192. doi:10.1186/s13046-022-02394-2.

Ciordia, S., Santos, F. M., Dias, J. M. L., Lamas, J. R., Paradela, A., Alvarez-Sola, G., et al. (2024). Refinement of paramagnetic bead-based digestion protocol for automatic sample preparation using an artificial neural network. Talanta 274. doi:10.1016/j.talanta.2024.125988.

Claveria-Cabello, A., Arechederra, M., Berasain, C., Fernández-Barrena, M. G., and Avila, M. A. (2021). Epigenetics in hepatoblastoma. Hepatoma Res. 7:65. doi:10.20517/2395-5079.2021.77.

Clavería-Cabello, A., Herranz, J. M., Latasa, M. U., Arechederra, M., Uriarte, I., Pineda-Lucena, A., et al. (2023). Identification and experimental validation of druggable epigenetic targets in hepatoblastoma. J. Hepatol. 79, 989–1005. doi:10.1016/J.JHEP.2023.05.031.

Colyn, L., Alvarez-Sola, G., Latasa, M. U., Uriarte, I., Herranz, J. M., Arechederra, M., et al. (2022). New molecular mechanisms in cholangiocarcinoma: signals triggering interleukin-6 production in tumor cells and KRAS co-opted epigenetic mediators driving metabolic reprogramming. J. Exp. Clin. Cancer Res. 41, 1–18. doi:10.1186/s13046-022-02386-2.

Colyn, L., Bárcena-Varela, M., Álvarez-Sola, G., Latasa, M. U., Uriarte, I., Santamaría, E., et al. (2021). Dual Targeting of G9a and DNA Methyltransferase-1 for the Treatment of Experimental Cholangiocarcinoma. Hepatology 73, 2380–2396. doi:10.1002/hep.31642.

DiPeri, T. P., Zhao, M., Evans, K. W., Varadarajan, K., Moss, T., Scott, S., et al. (2023). Convergent MAPK pathway alterations mediate acquired resistance to FGFR inhibitors in FGFR2 fusion-positive cholangiocarcinoma. J. Hepatol. doi:10.1016/J.JHEP.2023.10.041.

Dobin, A., Davis, C. A., Schlesinger, F., Drenkow, J., Zaleski, C., Jha, S., et al. (2013). STAR: Ultrafast universal RNA-seq aligner. Bioinformatics 29, 15–21. doi:10.1093/bioinformatics/bts635.

Dong, L., Lu, D., Chen, R., Lin, Y., Zhu, H., Zhang, Z., et al. (2022). Proteogenomic characterization identifies clinically relevant subgroups of intrahepatic cholangiocarcinoma. 40, 70–87.e15. doi:10.1016/J.CCELL.2021.12.006.

Dong, Z., Yuan, Z., Jin, T., Gao, C., Wang, X., and Xu, F. (2025). Lactate at the crossroads of tumor metabolism and immune escape: a new frontier in cancer therapy. J. Transl. Med. 23. doi:10.1186/S12967-025-07272-X.

Dragomir, M. P., Calina, T. G., Perez, E., Schallenberg, S., Chen, M., Albrecht, T., et al. (2023). DNA methylation-based classifier differentiates intrahepatic pancreato-biliary tumours. EBioMedicine 93. doi:10.1016/J.EBIOM.2023.104657.

Du, J., Johnson, L. M., Jacobsen, S. E., and Patel, D. J. (2015). DNA methylation pathways and their crosstalk with histone methylation. Nat. Rev. Mol. Cell Biol. 16, 519–32. doi:10.1038/nrm4043.

Ducker, G. S., and Rabinowitz, J. D. (2017). One-Carbon Metabolism in Health and Disease. Cell Metab. 25, 27–42. doi:10.1016/J.CMET.2016.08.009.

Elurbide, J., Colyn, L., Latasa, M. U., Uriarte, I., Mariani, S., Lopez-Pascual, A., et al. (2025). Identification of PRMT5 as a therapeutic target in cholangiocarcinoma. Gut 74, 116–127. doi:10.1136/gutjnl-2024-332998.

Esteve-Puig, R., Bueno-Costa, A., and Esteller, M. (2020). Writers, readers and erasers of RNA modifications in cancer. Cancer Lett. 474, 127–137. doi:10.1016/j.canlet.2020.01.021.

Feehley, T., O’Donnell, C. W., Mendlein, J., Karande, M., and McCauley, T. (2023). Drugging the epigenome in the age of precision medicine. Clin. Epigenetics 15. doi:10.1186/S13148-022-01419-Z.

Fernández-Barrena, M. G., Arechederra, M., Colyn, L., Berasain, C., and Avila, M. A. (2020). Epigenetics in hepatocellular carcinoma development and therapy: The tip of the iceberg. JHEP Reports 2, 100167. doi:10.1016/j.jhepr.2020.100167.

Fernandez-Barrena, M. G., Uriarte, I., Sarobe, P., and Avila, M. A. (2025). Epigenetic mechanisms in HCC immune landscape: Therapeutic implications. Semin. Immunol. 79, 101980. doi:10.1016/j.smim.2025.101980.

Feron, O. (2019). The many metabolic sources of acetyl-CoA to support histone acetylation and influence cancer progression. Ann. Transl. Med. 7, S277. doi:10.21037/atm.2019.11.140.

Ghisletti, S., and Russo, M. (2025). TCA-cycle metabolites in the nucleus: drivers of chromatin and epigenetic control. BMC Biol. 23, 316. doi:10.1186/s12915-025-02423-4.

Goeppert, B., Stichel, D., Toth, R., Fritzsche, S., Loeffler, M. A., Schlitter, A. M., et al. (2022). Integrative analysis reveals early and distinct genetic and epigenetic changes in intraductal papillary and tubulopapillary cholangiocarcinogenesis. Gut 71, 391–401. doi:10.1136/GUTJNL-2020-322983.

Guertin, D. A., and Wellen, K. E. (2023). Acetyl-CoA metabolism in cancer. Nat. Rev. Cancer 23, 156–172. doi:10.1038/s41568-022-00543-5.

He, J., Yao, W., Wang, J., Schemmer, P., Yang, Y., Liu, Y., et al. (2016). TACC3 overexpression in cholangiocarcinoma correlates with poor prognosis and is a potential anti-cancer molecular drug target for HDAC inhibitors. Oncotarget 7, 75441–75456. doi:10.18632/oncotarget.12254.

Heberle, H., Meirelles, V. G., da Silva, F. R., Telles, G. P., and Minghim, R. (2015). InteractiVenn: a web-based tool for the analysis of sets through Venn diagrams. BMC Bioinforma. 2015 *161* 16, 169-. doi:10.1186/S12859-015-0611-3.

Herranz, J. M., López-Pascual, A., Clavería-Cabello, A., Uriarte, I., Latasa, M. U., Irigaray-Miramon, A., et al. (2023). Comprehensive analysis of epigenetic and epitranscriptomic genes’ expression in human NAFLD. J. Physiol. Biochem. 79, 901–924. doi:10.1007/s13105-023-00976-y.

Hou, Y., Liu, W., Yi, X., Yang, Y., Su, D., Huang, W., et al. (2020). PHF20L1 as a H3K27me2 reader coordinates with transcriptional repressors to promote breast tumorigenesis. Sci. Adv. 6, eaaz0356. doi:10.1126/SCIADV.AAZ0356.

Hsiao, L. L., Dangond, F., Yoshida, T., Hong, R., Jensen, R. V., Misra, J., et al. (2001). A compendium of gene expression in normal human tissues. Physiol. Genomics 7, 97–104. doi:10.1152/PHYSIOLGENOMICS.00040.2001.

Hu, S., Molina, L., Tao, J., Liu, S., Hassan, M., Singh, S., et al. (2022). NOTCH-YAP1/TEAD-DNMT1 axis drives hepatocyte reprogramming into intrahepatic cholangiocarcinoma. Gastroenterology 163, 449. doi:10.1053/J.GASTRO.2022.05.007.

Huang, R., Wu, Y., and Zou, Z. (2022). Combining EZH2 inhibitors with other therapies for solid tumors: more choices for better effects. Epigenomics 14, 1449–1464. doi:10.2217/EPI-2022-0320.

Huo, M., Zhang, J., Huang, W., and Wang, Y. (2021). Interplay Among Metabolism, Epigenetic Modifications, and Gene Expression in Cancer. Front. cell Dev. Biol. 9, 793428. doi:10.3389/fcell.2021.793428.

Ilyas, S. I., Affo, S., Goyal, L., Lamarca, A., Sapisochin, G., Yang, J. D., et al. (2023). Cholangiocarcinoma - novel biological insights and therapeutic strategies. Nat. Rev. Clin. Oncol. 20, 470–486. doi:10.1038/S41571-023-00770-1.

Jo, C., Park, S., Oh, S., Choi, J., Kim, E. K., Youn, H. D., et al. (2020). Histone acylation marks respond to metabolic perturbations and enable cellular adaptation. Exp. Mol. Med. 2020 5212 52, 2005–2019. doi:10.1038/s12276-020-00539-x.

Job, S., Rapoud, D., Dos Santos, A., Gonzalez, P., Desterke, C., Pascal, G., et al. (2020). Identification of Four Immune Subtypes Characterized by Distinct Composition and Functions of Tumor Microenvironment in Intrahepatic Cholangiocarcinoma. Hepatology 72, 965. doi:10.1002/HEP.31092.

Johnson, W. E., Li, C., and Rabinovic, A. (2007). Adjusting batch effects in microarray expression data using empirical Bayes methods. Biostatistics 8, 118–127. doi:10.1093/biostatistics/kxj037.

Jung, D. E., Park, S. B., Kim, K., Kim, C., and Song, S. Y. (2017). CG200745, an HDAC inhibitor, induces anti-tumour effects in cholangiocarcinoma cell lines via miRNAs targeting the Hippo pathway. Sci. Rep. 7, 10921. doi:10.1038/s41598-017-11094-3.

Jusakul, A., Cutcutache, I., Yong, C. H., Lim, J. Q., Huang, M. N., Padmanabhan, N., et al. (2017). Whole-Genome and Epigenomic Landscapes of Etiologically Distinct Subtypes of Cholangiocarcinoma. Cancer Discov. 7, 1116–1135. doi:10.1158/2159-8290.CD-17-0368.

Kanehisa, M., Goto, S., Sato, Y., Kawashima, M., Furumichi, M., and Tanabe, M. (2014). Data, information, knowledge and principle: back to metabolism in KEGG. Nucleic Acids Res. 42, D199–205. doi:10.1093/nar/gkt1076.

Kechin, A., Boyarskikh, U., Kel, A., and Filipenko, M. (2017). CutPrimers: A New Tool for Accurate Cutting of Primers from Reads of Targeted Next Generation Sequencing. J. Comput. Biol. 24, 1138–1143. doi:10.1089/cmb.2017.0096.

Khan, M. A., Mishra, D., Kumar, R., and Siddique, H. R. (2025). Revisiting epigenetic regulation in cancer: Evolving trends and translational implications. Int. Rev. Cell Mol. Biol. 390, 1–24. doi:10.1016/bs.ircmb.2024.09.002.

Krieg, A. J., Rankin, E. B., Chan, D., Razorenova, O., Fernandez, S., and Giaccia, A. J. (2010). Regulation of the histone demethylase JMJD1A by hypoxia-inducible factor 1 alpha enhances hypoxic gene expression and tumor growth. Mol. Cell. Biol. 30, 344–53. doi:10.1128/MCB.00444-09.

Krill-Burger, J. M., Dempster, J. M., Borah, A. A., Paolella, B. R., Root, D. E., Golub, T. R., et al. (2023). Partial gene suppression improves identification of cancer vulnerabilities when CRISPR-Cas9 knockout is pan-lethal. Genome Biol. 24, 192. doi:10.1186/s13059-023-03020-w.

Lee, S., Lee, J., Chae, S., Moon, Y., Lee, H.-Y., Park, B., et al. (2017). Multi-dimensional histone methylations for coordinated regulation of gene expression under hypoxia. Nucleic Acids Res. 45, 11643–11657. doi:10.1093/nar/gkx747.

Leek, J. T., Johnson, W. E., Parker, H. S., Jaffe, A. E., and Storey, J. D. (2012). The SVA package for removing batch effects and other unwanted variation in high-throughput experiments. Bioinformatics 28, 882–883. doi:10.1093/bioinformatics/bts034.

Li, H., Peng, C., Zhu, C., Nie, S., Qian, X., Shi, Z., et al. (2021). Hypoxia promotes the metastasis of pancreatic cancer through regulating NOX4/KDM5A-mediated histone methylation modification changes in a HIF1A-independent manner. Clin. Epigenetics 13, 18. doi:10.1186/s13148-021-01016-6.

Li, X., Egervari, G., Wang, Y., Berger, S. L., and Lu, Z. (2018). Regulation of chromatin and gene expression by metabolic enzymes and metabolites. Nat. Rev. Mol. Cell Biol. 19, 563–578. doi:10.1038/s41580-018-0029-7.

Liang, Y., Bu, Q., You, W., Zhang, R., Xu, Z., Gan, X., et al. (2024). Single-cell analysis reveals hypoxia-induced immunosuppressive microenvironment in intrahepatic cholangiocarcinoma. Biochim. Biophys. acta. Mol. basis Dis. 1870, 167276. doi:10.1016/j.bbadis.2024.167276.

Liu, L., Zhen, X. T., Denton, E., Marsden, B. D., and Schapira, M. (2012). ChromoHub: A data hub for navigators of chromatin-mediated signalling. Bioinformatics 28, 2205–2206. doi:10.1093/bioinformatics/bts340.

Lori, G., Pastore, M., Navari, N., Piombanti, B., Booijink, R., Rovida, E., et al. (2024). Altered fatty acid metabolism rewires cholangiocarcinoma stemness features. JHEP Reports 6. doi:10.1016/j.jhepr.2024.101182.

Lu, Q., Ding, X., Huang, T., Zhang, S., Li, Y., Xu, L., et al. (2019). BRD4 degrader ARV-825 produces long-lasting loss of BRD4 protein and exhibits potent efficacy against cholangiocarcinoma cells. Am. J. Transl. Res. 11, 5728–5739. doi:https://pmc.ncbi.nlm.nih.gov/articles/PMC6789278.

Ma, W., Zhang, J., Chen, W., Liu, N., and Wu, T. (2024). Notch-Driven Cholangiocarcinogenesis Involves the Hippo Pathway Effector TAZ via METTL3-m6A-YTHDF1. Cell. Mol. Gastroenterol. Hepatol. 19, 101417. doi:10.1016/J.JCMGH.2024.101417.

Macias, R. I. R., Cardinale, V., Kendall, T. J., Avila, M. A., Guido, M., Coulouarn, C., et al. (2022). Clinical relevance of biomarkers in cholangiocarcinoma: critical revision and future directions. Gut 71, gutjnl-2022-327099. doi:10.1136/gutjnl-2022-327099.

Manzano-Núñez, F., Prates Tiago Aguilar, L., Sempoux, C., and Lemaigre, F. P. (2023). Biliary Tract Cancer: Molecular Biology of Precursor Lesions. Semin. Liver Dis. 43, 472–484. doi:10.1055/a-2207-9834.

Marakulina, D., Vorontsov, I. E., Kulakovskiy, I. V., Lennartsson, A., Drabløs, F., and Medvedeva, Y. A. (2023). EpiFactors 2022: expansion and enhancement of a curated database of human epigenetic factors and complexes. Nucleic Acids Res. 51, D564–D570. doi:10.1093/nar/gkac989.

Marin, J. J. G., Lozano, E., Herraez, E., Asensio, M., Di Giacomo, S., Romero, M. R., et al. (2018). Chemoresistance and chemosensitization in cholangiocarcinoma. Biochim. Biophys. acta. Mol. basis Dis. 1864, 1444–1453. doi:10.1016/j.bbadis.2017.06.005.

Martin-Serrano, M. A., Kepecs, B., Torres-Martin, M., Bramel, E. R., Haber, P. K., Merritt, E., et al. (2023). Novel microenvironment-based classification of intrahepatic cholangiocarcinoma with therapeutic implications. Gut 72, 736–748. doi:10.1136/GUTJNL-2021-326514.

Martin, M. (2011). Cutadapt removes adapter sequences from high-throughput sequencing reads. EMBnet.journal 17, 10. doi:10.14806/ej.17.1.200.

Martínez-Arranz, I., Mayo, R., Pérez-Cormenzana, M., Mincholé, I., Salazar, L., Alonso, C., et al. (2015). Enhancing metabolomics research through data mining. J. Proteomics 127, 275–288. doi:10.1016/j.jprot.2015.01.019.

Merino-Azpitarte, M., Lozano, E., Perugorria, M. J., Esparza-Baquer, A., Erice, O., Santos-Laso, Á., et al. (2017). SOX17 regulates cholangiocyte differentiation and acts as a tumor suppressor in cholangiocarcinoma. J. Hepatol. 67, 72–83. doi:10.1016/j.jhep.2017.02.017.

Meyers, R. M., Bryan, J. G., McFarland, J. M., Weir, B. A., Sizemore, A. E., Xu, H., et al. (2017). Computational correction of copy number effect improves specificity of CRISPR–Cas9 essentiality screens in cancer cells. Nat. Genet. 49, 1779–1784. doi:10.1038/ng.3984.

Momeni, K., Ghorbian, S., Ahmadpour, E., and Sharifi, R. (2023). Unraveling the complexity: understanding the deconvolutions of RNA-seq data. Transl. Med. Commun. 2023 81 8, 21-. doi:10.1186/S41231-023-00154-8.

Montal, R., Sia, D., Montironi, C., Leow, W. Q., Esteban-Fabró, R., Pinyol, R., et al. (2020). Molecular classification and therapeutic targets in extrahepatic cholangiocarcinoma. J. Hepatol. 73, 315–327. doi:10.1016/j.jhep.2020.03.008.

Morgan, M. A. J., and Shilatifard, A. (2020). Reevaluating the roles of histone-modifying enzymes and their associated chromatin modifications in transcriptional regulation. 52, 1271–1281. doi:10.1038/S41588-020-00736-4.

Natarajan, S. K., Pun, M., Haggerty-Skeans, J., and Venneti, S. (2025). Integration and Intersection of Cancer Metabolism with Epigenetic Pathways in Gliomas. Annu. Rev. Pathol. doi:10.1146/ANNUREV-PATHMECHDIS-111523-023424.

O’Rourke, C. J., Lafuente-Barquero, J., and Andersen, J. B. (2019). Epigenome Remodeling in Cholangiocarcinoma. Trends in cancer 5, 335–350. doi:10.1016/j.trecan.2019.05.002.

O’Rourke, C. J., Munoz-Garrido, P., Aguayo, E. L., and Andersen, J. B. (2018). Epigenome dysregulation in cholangiocarcinoma. Biochim. Biophys. acta. Mol. basis Dis. 1864, 1423–1434. doi:10.1016/j.bbadis.2017.06.014.

O’Rourke, C. J., Salati, M., Rae, C., Carpino, G., Leslie, H., Pea, A., et al. (2023). Molecular portraits of patients with intrahepatic cholangiocarcinoma who diverge as rapid progressors or long survivors on chemotherapy. Gut. doi:10.1136/gutjnl-2023-330748.

Oyon, D., Lopez-Pascual, A., Castello-Uribe, B., Uriarte, I., Orsi, G., Llorente, S., et al. (2025). Targeting of the G9a, DNMT1 and UHRF1 epigenetic complex as an effective strategy against pancreatic ductal adenocarcinoma. J. Exp. Clin. Cancer Res. 44, 13. doi:10.1186/s13046-024-03268-5.

Pajares, M. A., and Pérez-Sala, D. (2018). Mammalian Sulfur Amino Acid Metabolism: A Nexus Between Redox Regulation, Nutrition, Epigenetics, and Detoxification. Antioxid. Redox Signal. 29, 408–452. doi:10.1089/ars.2017.7237.

Pascale, A., Rosmorduc, O., and Duclos-Vallée, J.-C. (2023). New epidemiologic trends in cholangiocarcinoma. Clin. Res. Hepatol. Gastroenterol. 47, 102223. doi:10.1016/j.clinre.2023.102223.

Peng, L., Li, J., Wu, J., Xu, B., Wang, Z., Giamas, G., et al. (2021). A Pan-Cancer Analysis of SMARCA4 Alterations in Human Cancers. Front. Immunol. 12, 762598. doi:10.3389/fimmu.2021.762598.

Raggi, C., Taddei, M. L., Rae, C., Braconi, C., and Marra, F. (2022). Metabolic reprogramming in cholangiocarcinoma. J. Hepatol. 77, 849–864. doi:10.1016/j.jhep.2022.04.038.

Raggi, C., Taddei, M. L., Sacco, E., Navari, N., Correnti, M., Piombanti, B., et al. (2021). Mitochondrial oxidative metabolism contributes to a cancer stem cell phenotype in cholangiocarcinoma. J. Hepatol. 74, 1373–1385. doi:10.1016/j.jhep.2020.12.031.

Reina-Campos, M., Diaz-Meco, M. T., and Moscat, J. (2019). The complexity of the serine glycine one-carbon pathway in cancer. J. Cell Biol., jcb.201907022. doi:10.1083/jcb.201907022.

Robinson, M. D., McCarthy, D. J., and Smyth, G. K. (2009). edgeR: A Bioconductor package for differential expression analysis of digital gene expression data. Bioinformatics 26, 139–140. doi:10.1093/bioinformatics/btp616.

Rosario, S. R., Long, M. D., Affronti, H. C., Rowsam, A. M., Eng, K. H., and Smiraglia, D. J. (2018). Pan-cancer analysis of transcriptional metabolic dysregulation using The Cancer Genome Atlas. Nat. Commun. 9. doi:10.1038/s41467-018-07232-8.

Ruffolo, L. I., Jackson, K. M., Kuhlers, P. C., Dale, B. S., Figueroa Guilliani, N. M., Ullman, N. A., et al. (2022). GM-CSF drives myelopoiesis, recruitment and polarisation of tumour-associated macrophages in cholangiocarcinoma and systemic blockade facilitates antitumour immunity. Gut 71, 1386–1398. doi:10.1136/gutjnl-2021-324109.

Sarantis, P., Tzanetatou, E. D., Ioakeimidou, E., Vallilas, C., Androutsakos, T., Damaskos, C., et al. (2021). Cholangiocarcinoma: the role of genetic and epigenetic factors; current and prospective treatment with checkpoint inhibitors and immunotherapy. Am. J. Transl. Res. 13, 13246–13260. doi:https://pubmed.ncbi.nlm.nih.gov/35035673.

Satriano, L., Lewinska, M., Rodrigues, P. M., Banales, J. M., and Andersen, J. B. (2019). Metabolic rearrangements in primary liver cancers: cause and consequences. Nat. Rev. Gastroenterol. Hepatol. 16, 748–766. doi:10.1038/s41575-019-0217-8.

Sean, D., and Meltzer, P. S. (2007). GEOquery: A bridge between the Gene Expression Omnibus (GEO) and BioConductor. Bioinformatics 23, 1846–1847. doi:10.1093/bioinformatics/btm254.

Sia, D., Hoshida, Y., Villanueva, A., Roayaie, S., Ferrer, J., Tabak, B., et al. (2013). Integrative Molecular Analysis of Intrahepatic Cholangiocarcinoma Reveals 2 Classes That Have Different Outcomes. Gastroenterology 144, 829–840. doi:10.1053/j.gastro.2013.01.001.

Sin, S. Q., Mohan, C. D., Goh, R. M. W.-J., You, M., Nayak, S. C., Chen, L., et al. (2023). Hypoxia signaling in hepatocellular carcinoma: Challenges and therapeutic opportunities. Cancer Metastasis Rev. 42, 741–764. doi:10.1007/s10555-022-10071-1.

Sirica, A. E., Dumur, C. I., Campbell, D. J. W., Almenara, J. A., Ogunwobi, O. O., and Dewitt, J. L. (2009). Intrahepatic Cholangiocarcinoma Progression: Prognostic Factors and Basic Mechanisms. Clin. Gastroenterol. Hepatol. 7, S68–S78. doi:10.1016/j.cgh.2009.08.023.

Talom, A., Barhoi, A., Jirpu, T., Dawn, B., and Ghosh, A. (2025). Clinical progress and functional modalities of HDAC inhibitor-based combination therapies in cancer treatment. Clin. Transl. Oncol. doi:10.1007/S12094-025-03995-X.

Valle, J. W., Kelley, R. K., Nervi, B., Oh, D.-Y., and Zhu, A. X. (2021). Biliary tract cancer. *Lancet (London*, England) 397, 428–444. doi:10.1016/S0140-6736(21)00153-7.

Wang, Y. P., and Lei, Q. Y. (2018). Metabolic recoding of epigenetics in cancer. Cancer Commun. 2018 *381* 38, 25-. doi:10.1186/S40880-018-0302-3.

Wasenang, W., Puapairoj, A., Settasatian, C., Proungvitaya, S., and Limpaiboon, T. (2019). Overexpression of polycomb repressive complex 2 key components EZH2/SUZ12/EED as an unfavorable prognostic marker in cholangiocarcinoma. Pathol. Res. Pract. 215, 152451. doi:10.1016/j.prp.2019.152451.

Wu, G., Wang, Q., Wang, D., Xiong, F., Liu, W., Chen, J., et al. (2023). Targeting polycomb repressor complex 2-mediated bivalent promoter epigenetic silencing of secreted frizzled-related protein 1 inhibits cholangiocarcinoma progression. Clin. Transl. Med. 13, e1502. doi:10.1002/ctm2.1502.

Xiang, J., Luo, F., Chen, Y., Zhu, F., and Wang, J. (2014). Si-DNMT1 restore tumor suppressor genes expression through the reversal of DNA hypermethylation in cholangiocarcinoma. Clin. Res. Hepatol. Gastroenterol. 38, 181–189. doi:10.1016/j.clinre.2013.11.004.

Xie, M., Liang, H., Mao, Y., Yao, Y., and Tian, B. (2025). CBX3 Downregulates HLTF to Activate PI3K/AKT Signaling Promoting Cholangiocarcinoma. *Adv*. Biol. 9, e2400413. doi:10.1002/adbi.202400413.

Xiong, F., Wang, D., Xiong, W., Wang, X., Huang, W.-H., Wu, G.-H., et al. (2024). Unveiling the role of HP1α-HDAC1-STAT1 axis as a therapeutic target for HP1α-positive intrahepatic cholangiocarcinoma. J. Exp. Clin. Cancer Res. 43, 152. doi:10.1186/s13046-024-03070-3.

Xu, L., Yang, W., Che, J., Li, D., Wang, H., Li, Y., et al. (2022). Suppression of histone deacetylase 1 by JSL-1 attenuates the progression and metastasis of cholangiocarcinoma via the TPX2/Snail axis. Cell Death Dis. 13, 324. doi:10.1038/s41419-022-04571-9.

Yang, M., Soga, T., and Pollard, P. J. (2013). Oncometabolites: linking altered metabolism with cancer. J. Clin. Invest. 123, 3652–3658. doi:10.1172/JCI67228.

Yin, Y., Zhang, M., Dorfman, R. G., Li, Y., Zhao, Z., Pan, Y., et al. (2017). Histone deacetylase 3 overexpression in human cholangiocarcinoma and promotion of cell growth via apoptosis inhibition. Cell Death Dis. 8, e2856. doi:10.1038/cddis.2016.457.

Zach, S., Birgin, E., and Rückert, F. (2015). Primary cholangiocellular carcinoma cell lines. J. Stem Cell Res. Transplant. 2, 1013.

Zhang, C., Zhang, B., Meng, D., and Ge, C. (2019). Comprehensive analysis of DNA methylation and gene expression profiles in cholangiocarcinoma. Cancer Cell Int. 19. doi:10.1186/S12935-019-1080-Y.

Zhang, J., Chen, W., Ma, W., Han, C., Song, K., Kwon, H., et al. (2022). EZH2 Promotes Cholangiocarcinoma Development and Progression through Histone Methylation and microRNA-Mediated Down-Regulation of Tumor Suppressor Genes. Am. J. Pathol. 192, 1712–1724. doi:10.1016/j.ajpath.2022.08.008.

Zhang, Z.-J., Huang, Y.-P., Liu, Z.-T., Wang, Y.-X., Zhou, H., Hou, K.-X., et al. (2023). Identification of immune related gene signature for predicting prognosis of cholangiocarcinoma patients. Front. Immunol. 14, 1028404. doi:10.3389/fimmu.2023.1028404.

Zhao, M., Chen, X., Gao, G., Tao, L., and Wei, L. (2009). RLEdb: a database of rate-limiting enzymes and their regulation in human, rat, mouse, yeast and E. coli. Cell Res. 19, 793–5. doi:10.1038/cr.2009.61.

Zhao, R., Wang, N., Huang, H., Ma, W., and Yan, Q. (2014). CHD5 a tumour suppressor is epigenetically silenced in hepatocellular carcinoma. Liver Int. 34, e151–e160. doi:10.1111/LIV.12503;WGROUP:STRING:PUBLICATION.

Zhao, S., Allis, C. D., and Wang, G. G. (2021). The language of chromatin modification in human cancers. Nat. Rev. Cancer 21, 413–430. doi:10.1038/s41568-021-00357-x.

Zhong, B., Liao, Q., Wang, X., Wang, X., and Zhang, J. (2023). The roles of epigenetic regulation in cholangiocarcinogenesis. Biomed. Pharmacother. 166, 115290. doi:10.1016/j.biopha.2023.115290.

Zhong, Y.-J., Luo, X.-M., Liu, F., He, Z.-Q., Yang, S.-Q., Ma, W.-J., et al. (2024). Integrative analyses of bulk and single-cell transcriptomics reveals the infiltration and crosstalk of cancer-associated fibroblasts as a novel predictor for prognosis and microenvironment remodeling in intrahepatic cholangiocarcinoma. J. Transl. Med. 22, 422. doi:10.1186/s12967-024-05238-z.

Zhou, Y., Chen, Y., Zhang, X., Xu, Q., Wu, Z., Cao, X., et al. (2021). Brahma-Related Gene 1 Inhibition Prevents Liver Fibrosis and Cholangiocarcinoma by Attenuating Progenitor Expansion. Hepatology 74, 797–815. doi:10.1002/hep.31780.

