## Supplementary figures and images for "Transcriptomic profiling of epigenetic regulators and metabolic reprogramming in human cholangiocarcinoma"

### Supplementary Figure 1

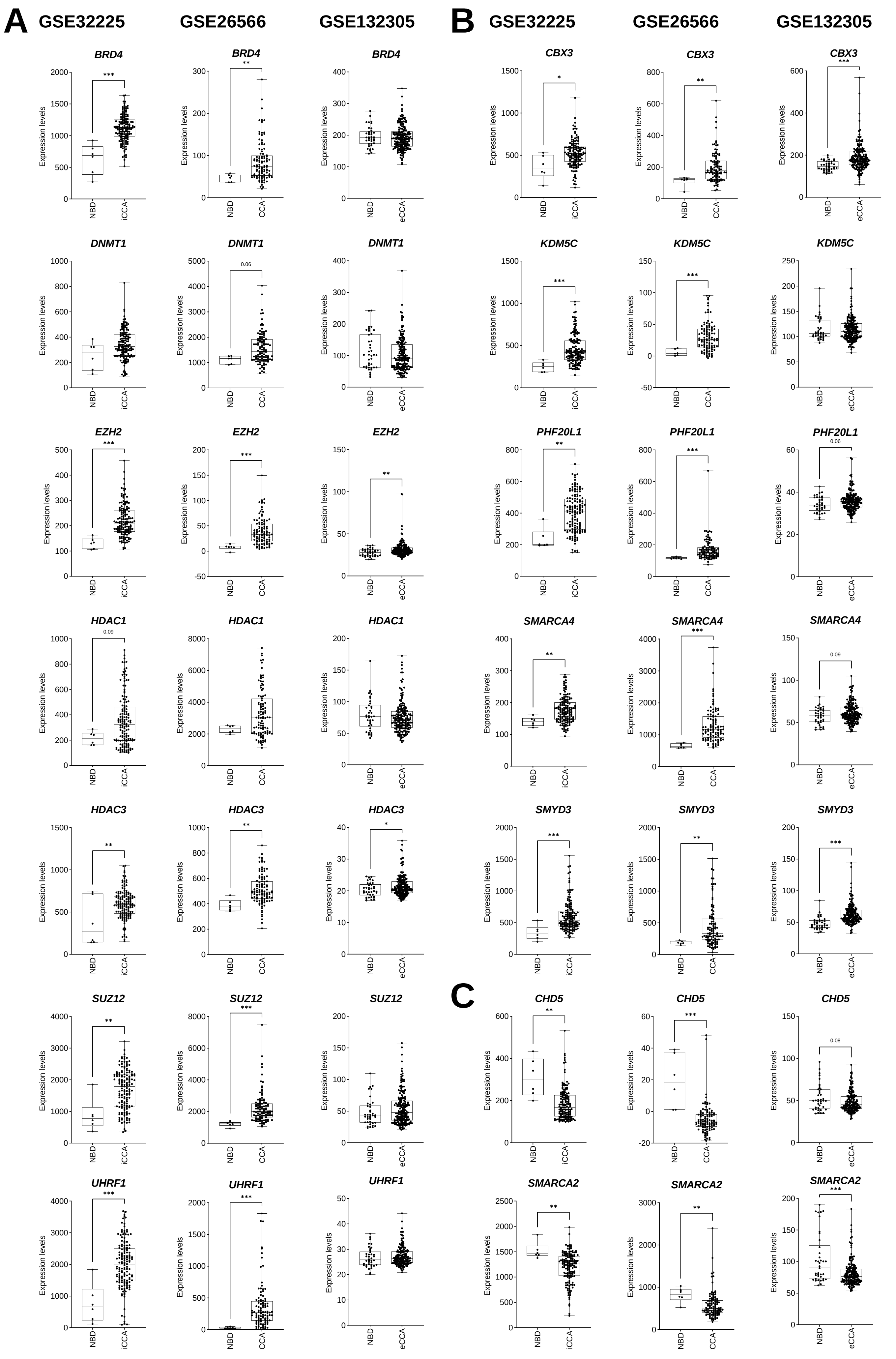

### Supplementary Figure 2

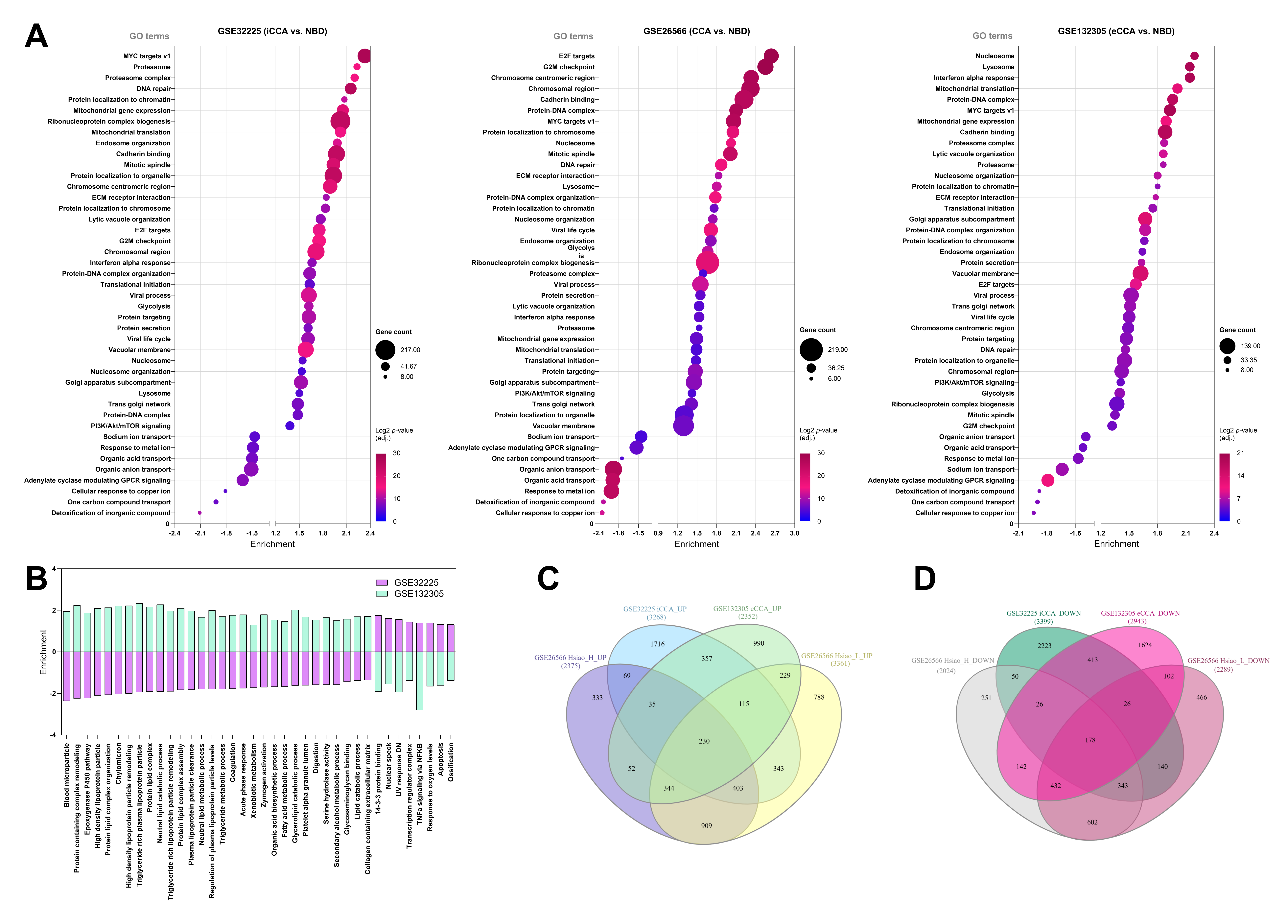

### Supplementary Figure 3

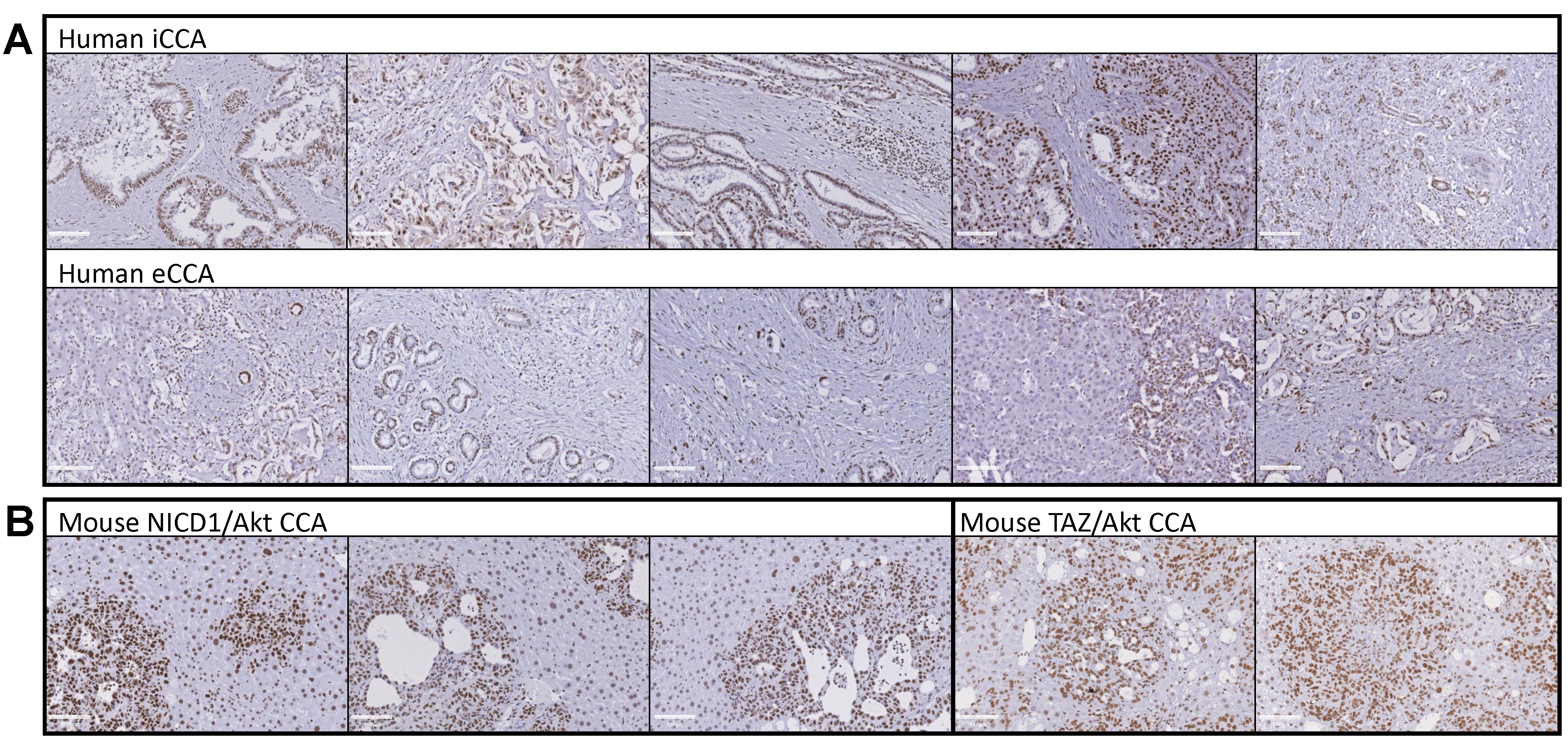
