## Supplementary Figures and Table legends for "Transcriptomic profiling of epigenetic regulators and metabolic reprogramming in human cholangiocarcinoma"

### *Supplementary Material*

#### 1 Supplementary Figures and Tables

**Supplementary Table 1.** List of epigenetic genes selected from the literature, classified by their epigenetic function and family, and manually curated functional activities.

**Supplementary Table 2.** List of metabolic genes functionally linked to epigenetic regulation, selected from the literature and classified according to the biological process in which they participate.5,10-CH+-THF, 5,10-Methylenetetrahydrofolate; DHF, Dihydrofolate; dTMP, Deoxythymidine monophosphate; fMet-tRNA, Formylmethionyl-tRNA; THF,Tetrahydrofolate; MTA, Methylthioadenosine; SAH, S-Adenosylhomocysteine; SAM, S-Adenosylmethionine; ACS-L/VL, Long- and very long-chain acyl-coA synthetase; ACS-S/M, Short- and medium-chain acyl-coA synthetase; CoA, Coenzyme A; PEP, Phosphoenolpyruvate; α-KG, Alpha-ketoglutarate.

**Supplementary Table 3.** List of KEGG metabolic pathways analyzed.

**Supplementary Table 4.** List of rate-limiting enzymes retrieved from RLEdb, and manually curated Enzyme Commission (EC) numbers, associated metabolic pathways, and functional classifications.

**Supplementary Table 5.** Intrahepatic cholangiocarcinoma (iCCA) patients' classification by transcriptomic signatures into the survival- and recurrence-related subclasses in the GSE32225.

**Supplementary Table 6.** Cholangiocarcinoma (CCA) patients' classification by transcriptomic signatures into the subclasses 1 and 2 and the Hsiao liver-specific subclasses in the GSE26566.

**Supplementary Table 7.** GO, KEGG, and Hallmark gene sets in GSEA analysis of differentially expressed genes between cholangiocarcinoma (CCA) and non-neoplastic bile duct epithelia (NBD) in GSE32225 (Sia et al., 2013), GSE26566 (Andersen et al., 2012), and GSE132305 (Montal et al., 2020). Enriched terms (over- or underrepresented) with statistically significant Normalized Enrichment Score (NES) (adjusted $p<0.05$) that included EpiGs among the DEGs were highlighted (≥10 highlighted in orange and ≥3 EpiGs in yellow).

**Supplementary Table 8.** Average expression ($Log_2FC$) of epigenetic genes (EpiGs), metabolic genes (MGs) and rate-limiting enzymes (RLEs) across GSE32225 (Sia et al., 2013), GSE26566 (Andersen et al., 2012), and GSE132305 (Montal et al., 2020) human datasets comparing cholangiocarcinoma (CCA) and non-neoplastic bile duct epithelia (NBD), human CCA tumoroids and healthy liver-derived organoids (Broutier et al., 2017), and TAZ/Akt-driven tumors and NICD1/Akt-driven tumors (early to advanced stages) compared to normal mouse livers.

**Supplementary Table 9.** GO, KEGG, and Hallmark gene sets in GSEA analysis of differentially expressed genes in cholangiocarcinoma (CCA) samples, comparing the worst- versus best-prognosis groups across the GSE32225 (Sia et al., 2013) and GSE26566 (Andersen et al., 2012) cohorts. Enriched terms (over- or underrepresented) with statistically significant Normalized Enrichment Score (NES) (adjusted $p<0.05$) that included EpiGs among the DEGs were highlighted (≥10 highlighted in orange and ≥3 EpiGs in yellow).

**Supplementary Table 10.** Differential expression of epigenetic genes (EpiGs), metabolic genes (MGs) and rate-limiting enzymes (RLEs) across tumor microenvironment- defined CCA immune-stromal clusters. Tumors were grouped into Immunogenic, Myeloid, Immune Desert, and Mesenchymal patterns using MCP-counter deconvolution of bulk transcriptomic data. Differentially expressed genes (raw $p<0.05$) were identified in GSE32225 (Sia et al., 2013) and Hsiao liver-specific subclasses in the GSE26566.

#### 1.1 Supplementary Figures

**Supplementary Figure 1. Expression of selected EpiGs in human CCA. (A)** Epigenetic genes (EpiGs) that have been reported as overexpressed in cholangiocarcinoma (CCA). **(B)** Additional upregulated and **(C)** downregulated EpiGs with limited or no prior evidence in CCA, representing potentially novel candidates.

**Supplementary Figure 2. Shared and divergent pathway enrichment and gene expression in human iCCA and eCCA. (A)** Subset of pathways consistently altered in cholangiocarcinoma (CCA) compared with normal bile duct (NBD) samples across GSE32225, GSE26566, and GSE132305 human datasets. **(B)** Pathways showing opposite enrichment directions between iCCA (GSE32225) and eCCA (GSE132305) compared with normal bile duct (NBD) samples. Overlapping differentially expressed genes (DEGs) across all comparisons, highlighting a subset of consistently **(C)** upregulated and **(D)** downregulated genes, including EpiGs, MGs, and RLEs, reflecting shared transcriptional programs in CCA.

**Supplementary Figure 3. SMARCA4 protein expression in human and experimental CCA. (A)** Representative SMARCA4 immunostaining in human intrahepatic and extrahepatic cholangiocarcinoma (iCCA and eCCA) samples and in **(B)** TAZ/Akt and NICD1/Akt mouse CCA models. Scale bar: 100 µm.
